# BRAF/MEK Inhibition Induces Cell State Transitions Boosting Immune Checkpoint Sensitivity in BRAF^V600E^-mutant Glioma

**DOI:** 10.1101/2023.02.03.526065

**Authors:** Yao Lulu Xing, Dena Panovska, Jong-Whi Park, Stefan Grossauer, Katharina Koeck, Brandon Bui, Emon Nasajpour, Jeffrey J. Nirschl, Zhi-Ping Feng, Pierre Cheung, Pardes Habib, Ruolun Wei, Jie Wang, Wes Thomason, Joanne Xiu, Alexander Beck, Katharina J. Weber, Patrick N. Harter, Michael Lim, Kelly Mahaney, Laura M. Prolo, Gerald A. Grant, Xuhuai Ji, Kyle M. Walsh, Jean M. Mulcahy Levy, Dolores Hambardzumyan, Claudia K. Petritsch

## Abstract

Resistance to BRAF plus MEK inhibition (BRAFi+MEKi) in BRAF^V600E^-mutant gliomas drives rebound, progression, and high mortality, yet it remains poorly understood. This study addresses the urgent need to develop treatments for BRAFi+MEKi-resistant glioma in novel mouse models and patient-derived materials. BRAFi+MEKi reveals glioma plasticity by heightening cell state transitions along glial differentiation trajectories, giving rise to astrocyte- and immunomodulatory oligodendrocyte (OL)-like states. PD-L1 upregulation in OL-like cells links cell state transitions to tumor evasion, possibly orchestrated by Galectin-3. BRAFi+MEKi induces interferon response signatures, tumor infiltration, and suppression of T cells. Combining BRAFi+MEKi with immune checkpoint inhibition enhances survival in a T cell-dependent manner, reinvigorates T cells, and outperforms individual or sequential therapies in mice. Elevated PD-L1 expression in BRAF-mutant versus BRAF-wildtype glioblastoma supports the rationale for PD-1 inhibition in patients. These findings underscore the potential of targeting glioma plasticity and highlight combination strategies to overcome therapy resistance in BRAF^V600E^-mutant HGG.

**In brief:** Xing et al. show that combined BRAF and MEK inhibitor (BRAFi+MEKi) treatment induces cell state transitions in BRAF^V600E^-mutant high-grade glioma cells linked with programmed death-ligand (PD-L1) upregulation and T cell suppression, potentially orchestrated through the secretion of galectin-3. These tumor-intrinsic adaptations may be overcome by concurrent immune checkpoint inhibition (ICI), as demonstrated in murine studies, offering novel therapeutic opportunities.

*Highlights:*

- BRAF^V600E^-mutant HGG exhibits cell plasticity induced by BRAFi+MEKi, which links cell state transitions towards glial differentiation with immune evasion
- BRAFi+MEKi enhances anti-tumor immunity and simultaneously suppresses T cells via PD-L1 upregulation
- BRAF-mutant glioblastoma has elevated PD-L1 expression compared to BRAF-wildtype counterparts, providing a criterion for PD-1 inhibition therapy
- Concurrent BRAFi+MEKi and immune checkpoint inhibition enhance T cell-mediated anti-tumor activity and boost survival more effectively than sequential treatment in mice, guiding clinical translation

## INTRODUCTION

Brain tumors cause the most years of life lost among cancers and are the leading cause of cancer-related death in children. ^1–4^ Unlike other cancers, molecular targeted therapies for brain tumors remain limited, with notable exceptions including inhibitors of BRAF (v-raf murine sarcoma viral oncogene homolog B1) and IDH (isocitrate dehydrogenase), now in clinical use for the treatment of patients with BRAF- and IDH-mutant gliomas, respectively. ^5,6^ BRAF, a kinase in the mitogen-activated protein kinase (MAPK) pathway, links growth factor signals to transcriptional changes, thereby orchestrating cell proliferation, survival, and differentiation.^7^ It is the most mutated kinase in cancer, with the V600E mutation comprising 90% of BRAF alterations.^8^ BRAF^V600E^ occurs in 7% of pediatric and 4% of adult primary brain tumors, with high prevalence in epithelioid glioblastoma (GBM), pleomorphic xanthoastrocytoma, and ganglioglioma. ^9–11^ Particularly common in pediatric low-grade gliomas (pLGG), BRAF^V600E^ confers poor outcomes with conventional therapies and increases risk of malignant progression to high-grade gliomas (HGG), especially with CDKN2A deletion. ^10,12–15^ The BRAF^V600E^ mutation generates a BRAF kinase monomer, which constitutively activates MAPK signaling and disables its feedback inhibition, promoting transcriptional output and cellular transformation. ^7,16,17^

ATP-competitive BRAF monomer inhibitors (BRAFi) – vemurafenib, dabrafenib, and encorafenib – effective in melanoma^18–22^, were repurposed to expand treatment options for patients with BRAF^V600E^-mutant glioma alongside broader screening for BRAF^V600E^ as a biomarker. ^7^ However, these BRAF inhibitors show limited efficacy as monotherapy, due to multiple adaptive resistance mechanisms, collectively known as the RAF inhibitor paradox. ^7^ As clinical benefit reportedly correlates with sustained MAPK suppression, ^23^ strong preclinical and clinical data support combining BRAF with MEK inhibitors (MEKi), targeting the downstream MAPK kinase to enhance treatment durability in HGG. ^24–26^ Combined, concurrent BRAF plus MEK inhibition (BRAFi+MEKi) outperforms chemoradiation in pLGG and shows encouraging response rates (∼47% in pLGG, ∼54% in adult LGG, and ∼33% in adult HGG) and reduced toxicities of combination therapy versus monotherapy. ^27–29^ In 2022, dabrafenib plus the allosteric MEKi trametinib received histology-agnostic, accelerated U.S. FDA approval for patients with unresectable or metastatic BRAF^V600E/K^-mutant solid tumors who have progressed following prior treatment and lack satisfactory alternatives. ^6,30^ In 2023, this combination was also approved for pediatric patients with BRAF^V600E^-mutant LGG who require systemic therapy. ^6,31^

Despite promising objective response rates (ORRs), most BRAF^V600E^-mutant gliomas resist BRAFi+MEKi, with residual tumor driving rebound, unbridled progression, and mortality. ^7,27,28,32–38^ Resistance mechanisms to BRAFi+MEKi are well-characterized, particularly in melanoma, but remain understudied in gliomas. Nevertheless, both tumor types exhibit intrinsic, acquired, and adaptive resistance to BRAFi. Shared resistance mechanisms include MAPK reactivation (e.g., via receptor tyrosine kinase activation), ^7,32,35,39^ and MAPK-independent mechanisms like PI3K/AKT/mTORC1 activation. ^26,32,39,40^ Unlike melanoma, BRAF^V600E^-mutant HGGs exhibit a high level of intrinsic (primary) resistance due to concomitant genetic alterations,^36^ including CDKN2A deletion and PI3K-mTOR mutations. ^41,42^ In addition, high intratumoral heterogeneity could be responsible for intrinsic resistance, with a polo-like-kinase 1 (PLK1)-high subpopulation characterized by prolonged G2-M phase, increased asymmetric cell division, and BRAFi+MEKi resistance, driving recurrence. ^43^ Hence, preclinical studies to enhance BRAFi efficacy involve combining BRAF^V600E^ inhibitors with PLK1, MEK, mTOR, or EGFR inhibitors. ^39,43–47^ However, only a few combinations have advanced to clinical trials (NCT04201457, NCT03919071).

Furthermore, intrinsic resistance could be attributed to the highly immunosuppressive glioma tumor microenvironment (TME), making immunotherapy for gliomas extremely challenging, in contrast to melanoma’s immunologically ‘hot’ TME characterized by robust T cell infiltration. ^48–51^ Programmed cell death protein-1 (PD-1) and cytotoxic T-lymphocyte antigen-4 (CTLA-4) immune checkpoints, exploited by tumors to evade immunity, offer therapeutic targets for immune checkpoint inhibitors (ICIs) ^52–54^ and were effective in BRAF^V600E^-mutant metastatic melanoma. ^55,56^ However, ICIs have demonstrated limited efficacy in HGGs. ^57,58^ Encouraging responses in replication-repair-deficient HGG and evidence of immune synergism with PD-1 blockade and MEKi (trametinib) in a small number of patients have been reported.^59^ Moreover, ERK1/2 phosphorylation predicts survival following anti-PD-1 immunotherapy in recurrent GBM.^60^

Considering the variability in BRAFi+MEKi responses across tumor lineages, ^36^ and the distinct intra- and inter-tumoral heterogeneity and TME in BRAF^V600E^-mutant HGGs, their responses to BRAFi+MEKi treatment require deeper investigation. ^48–51^ Moreover, growing evidence suggests that therapies tap into tumor plasticity, forcing cells to transition to distinct states. These cell states enable tumor cells to adapt to treatment and survive, ultimately driving resistance in solid cancers. ^61^ Given the widespread clinical use of BRAFi+MEKi and the high resistance rates to these inhibitors in HGGs, understanding treatment adaptations is critical. Due to limited patient cohorts, particularly with BRAFi+MEKi treatment, cell state transitions in response to treatment in BRAF^V600E^-mutant glioma are unknown. Robust non-human models (e.g., mice) are essential tools to investigate potential glioma cell adaptations and cell plasticity in BRAF^V600E^-mutant HGGs.

In this study, we investigated changes in HGG cell states in response to BRAFi+MEKi and understand cell plasticity as a potential adaptive resistance mechanism of BRAF^V600E^-mutant HGG. Pre- and post-BRAFi+MEKi-treatment patient samples, two patient-derived cell lines, and two novel syngeneic murine models of BRAF^V600E^-mutated HGG with varied treatment responsiveness were analyzed. BRAFi+MEKi heightened cell state transitions along glial differentiation trajectories, giving rise to astrocyte (AC)- and oligodendrocyte (OL)-like states. PD-L1 upregulation in OL-like cells linked cell state transitions to tumor evasion, possibly orchestrated by Galectin-3. BRAFi+MEKi-induced adaptive changes to the transcriptome included upregulation of IFNγ response gene expression, with antigen presentation programs, suggesting enhanced T cell-mediated anti-tumor immunity and potentially linking elevated cytotoxicity and antigen presentation to therapy response. Therapy-resistant murine tumors exhibited diminished cytotoxicity and impaired antigen presentation to T cells, suggesting immune evasion. This effect can be reversed by concurrently combining BRAFi+MEKi with ICI treatment, leading to significantly improved survival in a T cell-dependent manner. PD-L1 levels are also consistently elevated in BRAF-mutant versus BRAF-wildtype GBM patient cohorts, suggesting a pervasive immune evasion mechanism. Collectively, our parallel investigations of patient tumor samples, high-fidelity murine models, and patient-derived cell lines reveal cell plasticity and offer insights into BRAFi+MEKi modulation of tumor cell states and the tumor immune microenvironment, and may hold promise for a rational strategy combining BRAFi+MEKi with ICIs to improve outcomes in BRAF^V600E^-mutant HGGs.

## RESULTS

### BRAF^V600E^-mutant HGG Undergo Cell State Transitions Along Glial Differentiation Trajectories Following Extensive Treatment in Patients

Cancer treatment induces cellular adaptations via non-genetic mechanisms to drive resistance. Adaptive responses to BRAFi+MEKi in HGG are poorly understood. To investigate potential cellular adaptations to BRAFi+MEKi treatment, we analyzed the transcriptomes of paired BRAF^V600E^-mutant HGG from patients before dabrafenib+trametinib treatment and at recurrence using gene ontology (GO) analysis of bulk RNA sequencing (RNA-seq). Magnetic resonance imaging (MRI) confirmed recurrence, and immunofluorescence (IF), and immunohistochemistry (IHC) were done for validation **(Figures 1A-F and S1, Table S1)**. GO analyses showed upregulation of genes engaged in gliogenesis, synapse organization, axonogenesis, and cell-cell adhesion amongst other pathways after BRAFi+MEKi treatment **(Figure 1C and S1D).**

**Figure 1.**
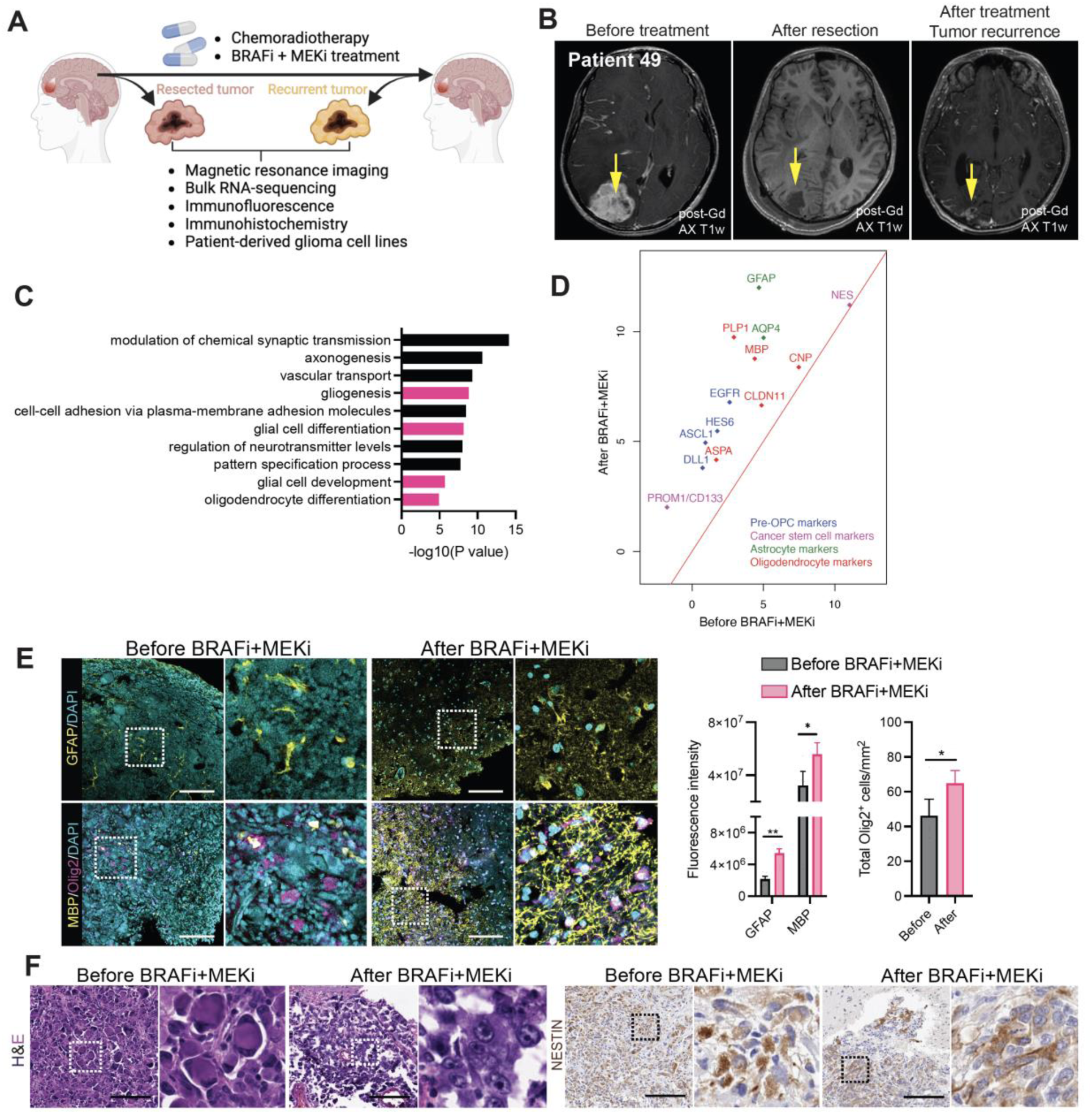
BRAF^V600E^-mutant HGG Undergo Cell State Transitions Along Glial Differentiation Trajectories Following Extensive Treatment in Patients. (A) Schematic of longitudinal analyses of BRAF^V600E^-mutant GBM from two patients. (B) Contrast-enhanced axial T1 MRI showing tumor (left), post-surgical cavity (middle), and recurrence (right) after BRAFi+MEKi in patient 49. (C) Bar plots of gliogenesis-related pathways (pink) from GO analysis at second recurrence after BRAFi+MEKi in patient 12. (D) Scatter plot of differentially expressed glial- and progenitor-related transcripts before and after BRAFi+MEKi in patient 12. (E) Representative IF images for GFAP, MBP, and OLIG2 before and after BRAFi+MEKi and chemoradiation in patient 49. Scale bars: 180µm (left), 140µm (right). Quantification of GFAP and MBP intensities and OLIG2+ cell frequencies in patient 49 (5 images of the tumor field were analyzed). High-magnification images of the white-boxed areas are shown on the right. *P<0.05 and **P<0.01 (Student’s t-test). Mean±SEM. (F) Representative H&E and NESTIN-stained tumor sections before and after BRAFi+MEKi in patient 49. High-magnification images of boxed areas are shown on the right. Scale bars: 150µm.

We further investigated pathways associated with gliogenesis for specific cell states and found that they are significantly enriched for markers for neural stem/progenitor cell (NSPCs) (*PROM1/CD133*), pre-oligodendrocyte progenitor cell (pre-OPCs)(*ASCL1, HES6, EGFR, DLL1*), astrocyte (AC) (*GFAP, AQP4),* myelin-forming oligodendrocyte (OL) (*PROM1/CD133, MBP, PLP1*)^62^ and -albeit at lower rates - mature OL (*ASPA, CNP, CLDN11*) states **(Figures 1D and S1D-E)**. Increased PROM1/CD133+ expression aligns with previous findings showing a CD133+ BRAF inhibitor-resistant cell population in BRAF^V600E^-mutant GBM.^43^ PROM1/CD133 and NESTIN expression – typically associated with a stemness state in GBM – were not co-regulated since NESTIN transcript **(Figure 1D)** and protein levels **(Figure 1F)** were unchanged after BRAFi+MEKi treatment. Further IF validation confirmed the upregulation of selected glial markers at the protein level, showing increased GFAP, MBP, and OLIG2 expression in post-treatment samples **(Figure 1E)**.

In conclusion, heavily treated BRAF^V600E^-mutant HGG retain NSPC markers and transition to AC-, pre-OPC-, and OL-like states, the latter reminiscent of pre-myelinating and mature OL-like cells.

### Novel Orthotopic BRAF^V600E^-mutant Mouse Models Show Intra-tumoral Cellular Heterogeneity

To address small patient cohort limitations of BRAF^V600E^-mutant HGG, particularly for BRAFi+MEKi treatment, we developed novel mouse models. We generated two BRAF^600E^-mutant HGG models (BRAF-M34 and RCAS-BRAF), with distinct co-mutations on an identical genetic background (C57BL/6), inducing gliomas at 54 days and 70 days post-injection, respectively **(Figures 2A, 2F and S2A-B, Table S2, STAR Methods)**. We subsequently isolated cells from these endogenous BRAF-M34 and RCAS-BRAF HGGs, and re-injected dissociated cells into C57BL/6 mouse brains to generate large orthotopic masses with high tumor take rate (BRAF-M34: 95%; RCAS-BRAF: 90%) **(Figure 2A, Table S2)**, as visualized by MRI **(Figures 2B, 2G)**. Histopathologic analyses of orthotopic masses revealed HGG features, including high nuclear pleomorphism, necrosis, and microvascular proliferation, with subtle model-specific differences **(Figures 2C, 2H, S2C and S2I, Table S2, STAR Methods)**. IF revealed high intra-tumoral heterogeneity in BRAF-M34 and RCAS-BRAF orthotopic glioma tested alongside BRAF-2341, which we previously generated on the FVB/N background.^26^ Expression of GFAP, OLIG2, NESTIN, PROM1/CD133, and OPC markers PDGFRα and NG2 in single tumor entities were detected **(Figures 2D-E, 2I, S2C-H, S2J, Table S2)**. Glial and neuronal markers are no longer lineage-restricted as in the normal brain but frequently co-expressed in orthotopic HGG, consistent with human HGGs **(Figure 2I)**. ^63^ Reactive ACs (GFAP+) and glioma-associated OPCs (PDGFRα/NG2) and OLs (nuclear OLIG2/PROM1), ^62,64^ identified by lack of GFP and CRE expression, were found in all three models as part of the HGG TME **(Figures 2D, middle panel, S2C-D, S2H)**. PROM1/CD133 was co-expressed with NESTIN, potentially marking an HGG cell state associated with stemness **(Figure 2E)**, and with OLIG2, which encompasses highly-proliferative OPC-like states in addition to the non-proliferative, mature OLs **(Figures S2F, S2H, S2J)** and - when cytoplasmic - also ACs **(Figure S2J)**. ^65^ Staining with proliferation marker Ki67 revealed that PROM1/CD133-positive cells are less proliferative than OLIG2-single-positive cells, suggesting that the former cell population has a cell state consistent with differentiation into OL-like cells. ^62^ Our two novel murine models (BRAF-M34, RCAS-BRAF) recapitulate features of human BRAF^V600E^-mutant HGG, ^66^ including intra-tumoral cellular heterogeneity, with model-specific differences. These models provide robust preclinical tools, facilitating immunological studies due to their intact immune system and cross-model comparisons owing to their identical C57BL/6 genetic backgrounds.

**Figure 2.**
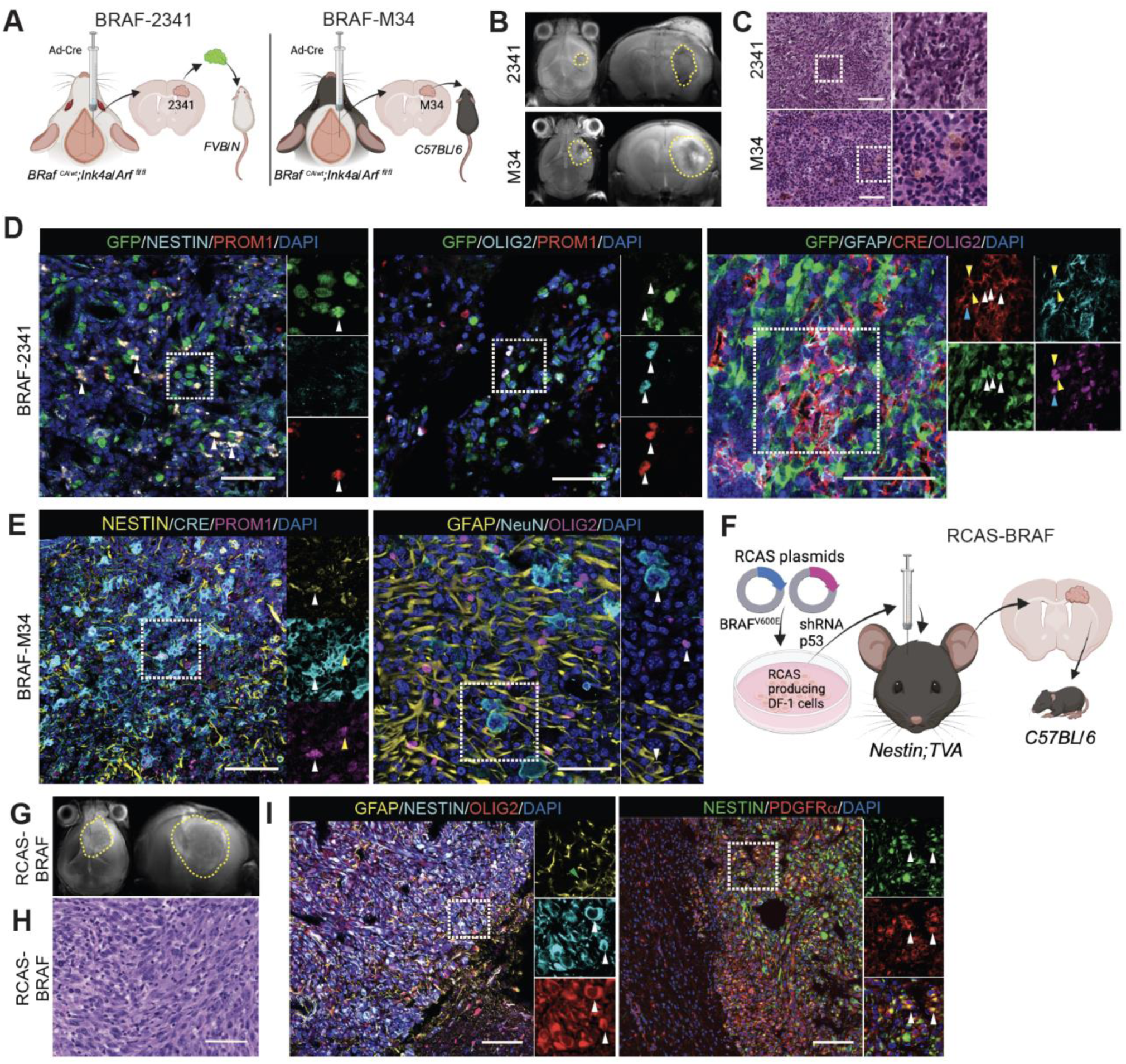
Novel Orthotopic BRAF^V600E^-mutant Mouse Models Show Intra-tumoral Cellular Heterogeneity. (A) Schematic diagram depicting the generation of BRAF-2341 and BRAF-M34 models and generation of the orthotopic, immunocompetent models syngeneic with FVB/N and C57BL/6 backgrounds, respectively. BRAF-2341 HGG cells were modified with lentivirus encoding GFP before orthotopic injection. (B) Representative Z-projection of T2-weighted MRI scans in both axial and coronal orientations with yellow dashed lines outlining the tumor in the injected right hemisphere of mice with BRAF-2341 and BRAF-M34 BRAF^V600E^-mutant, CDKN2A-deleted HGGs. (C) Representative low and high magnification images of H&E-stained coronal sections of BRAF-2341 HGG and BRAF-M34 HGG. Scale bars: 80µm (top), 60µm (bottom) **(D)** Representative IF images of BRAF-2341. GFP and CRE expression marks tumor cells. DAPI (cell nuclei). Left panel: GFP and PROM1/CD133 double-positive tumor cells (white arrowheads). Middle panel: OLIG2 and PROM1/CD133 double-positive/GFP-negative glioma-associated OLs (white arrowheads). Right panel: GFP and CRE double-positive tumor cells (white arrowheads); tumor cells co-express GFAP and OLIG2 (yellow arrowheads); or OLIG2 alone (blue arrowhead). Single-channel images of the white-boxed area are shown next to the merged images. Scale bars: 60µm (left), 45µm (middle), 75µm (right). **(E)** Representative IF images of BRAF-M34 HGG. Left panel: CRE, NESTIN, and PROM1/CD133 triple-positive NSPC/CSC-like tumor cell (white arrowheads). CRE and PROM1/CD133 double-positive (NESTIN negative) OL-like tumor cells (yellow arrowheads). Right panel: NeuN, OLIG2, and GFAP single positive cells (white arrowheads) in the tumor core featuring hypercellularity. Single-channel images of the white-boxed area are shown next to the merged images. Scale bars: 110µm (left), 50µm (right). (F) Schematic diagram describing the generation of RCAS-induced BRAF^V600E^ TP53-deleted (RCAS-BRAF) HGG and orthotopic, immunocompetent model development in Nestin-Tva mice syngeneic with C57BL/6. (G) Representative Z-projection of T2-weighted MRI scans in both axial and coronal orientations with yellow dashed outlines of the tumor infiltrating into the contralateral hemisphere. (H) Representative image of an H&E-stained coronal section of orthotopic RCAS-BRAF HGG showing hypercellular proliferation of malignant ovoid to spindled cells arranged in fascicles and sheets. Scale bar: 140µm. (I) Representative IF images of RCAS-BRAF HGG. GFAP-positive astrocyte-like cells, infrequently double-positive for OLIG2 (green arrowhead). An abundance of NESTIN+ cells, frequently double-positive for OLIG2 (white arrowheads), and PDGFRα+ cells (white arrowheads). Single-channel images of the white-boxed area are shown next to the merged images. Scale bars: 90µm (left), 115µm (right).

### BRAFi+MEKi Treatment Heightens Cell State Transitions Along Glial Differentiation Trajectories in BRAF^V600E^-mutant HGG

Our analyses of heavily pretreated BRAF^V600E^-mutant HGG patient materials strongly suggested that BRAFi+MEKi treatment boosts cell states associated with glial differentiation **(Figures 1D and S1D)**. To investigate these changes in additional samples, we assessed whether BRAFi+MEKi treatment boosts similar cell state changes in murine BRAF^V600E^-mutant HGG. Moreover, we used single-cell RNA sequencing (scRNA-seq) of tumors in addition to bulk RNA-seq to analyze these changes on a more granular level. Established (BRAF-2341) and novel (RCAS-BRAF) mouse models were treated with dabrafenib+trametinib for up to 14 days as indicated, and tumors were analyzed by bulk RNA-seq or scRNA-seq, then flow cytometry (FC) and IF for validation **(Figures 3A and 3C)**. In BRAF-2341 HGG cells, isolated by FC sorting for A2B5, ^67^ BRAFi+MEKi boosted the expression of transcripts associated with myelin-forming OL-(*Plp1, Mbp, Cnp, Prom1/CD133, Cldn11*), mature OL- (*Aspa*), and AC-like states (*Gfap, Clu*) **(Figures 3B and S1E)**. We confirmed that BRAFi+MEKi upregulated transcripts associated with AC- and OL-like states in two patient-derived BRAF^V600E^-mutant HGG cell lines (aGBM5 and STN-49), albeit with quantitative differences in certain transcripts **(Figures S3A-B)**, and consistent with their selected upregulation in patient tumors **(Figures 1D-E)**. Additionally, BRAFi+MEKi boosted the expression of transcripts associated with myelin-forming OL- (*Mbp, Plp1, Pllp, Prom1*) and AC-like states (*Gfap, Aqp4*) in the RCAS-BRAF model, as analyzed by scRNA-seq and validated by IF **(Figures 3C, S1E, S3F-G)**. GO analysis revealed significant upregulation of transcripts involved in gliogenesis **(Figure 3E)**, while IF confirmed upregulation of GFAP and MBP proteins, associated with glial differentiation (AC and OL), respectively **(Figures 3F-G)**. Markers for NSPC- *(Nes)*, pre-OPC-*(Dll1, Egfr, Hes6)*, and OPC-like (*Pdgfrα, Cspg4*) states were downregulated by BRAFi+MEKi treatment in BRAF-2341 cells **(Figures 3B and S1E)**, suggesting that cells with NSPC-, pre-OPC-, and OPC-like states are sensitive to short-term 3-day treatment. Decreased expression of transcripts for NSPC- and OPC-like states (*Nes*, *Cspg4*) was also detected, whereas some markers for pre-OPC-like states *(Ascl1, Hes6, Egfr)* were unchanged in the RCAS-BRAF tumors **(Figure 3D)** and OPC-like state marker PDGFRα protein was reduced, as shown by IF **(Figure 3G)**.

**Figure 3.**
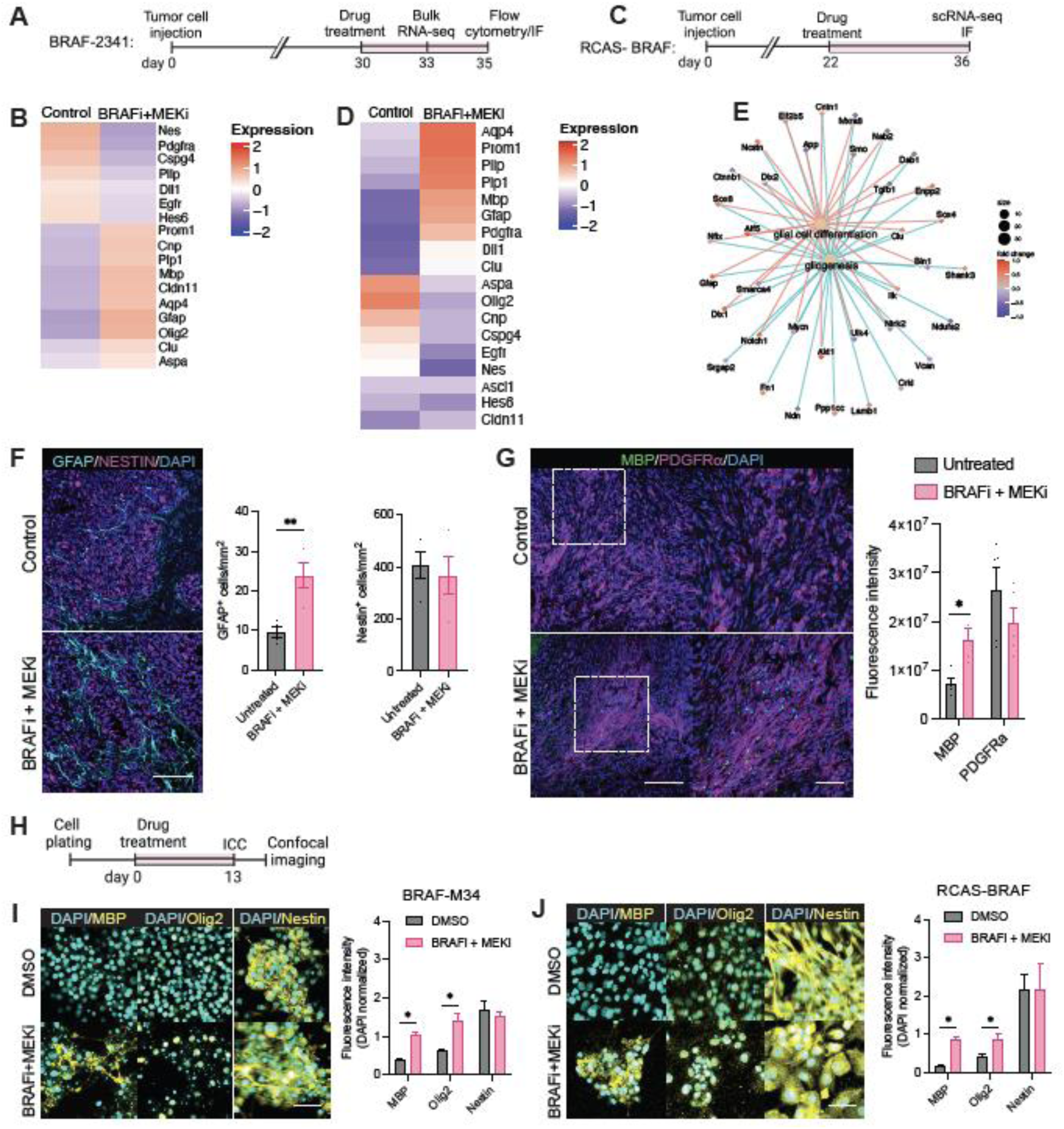
BRAFi+MEKi Treatment Heightens Cell State Transitions Along Glial Differentiation Trajectories in BRAF^V600E^-mutant HGG. (A) Schematic for experimental timeline for BRAFi+MEKi treatment and analyses (RNA-seq, flow cytometry=FC, and immunofluorescence=IF) in the BRAF-2341 model. (B) Heatmap from bulk RNA-seq depicting average expression levels of stem/progenitor cell and glial differentiation-associated transcripts between two replicates in BRAF-2341 HGG after 3-day BRAFi+MEKi treatment. (C) Schematic for experimental timeline for 14-day BRAFi+MEKi treatment and analyses (scRNA-seq, IF) of RCAS-BRAF HGG. (D) Heatmap from scRNA-seq data showing transcripts in the tumor cluster. Transcripts associated with myelin-forming OL- (*Mbp, Plp*) and AC-like (*Aqp4, Gfap, Clu*) cell states are upregulated, while OPC (*Cspg4*) and NSPC/CSC markers (*Nes*) are downregulated. (E) Cnetplot showing enrichment of glial-related transcripts from GO analysis of the scRNA-seq dataset from RCAS-BRAF HGG. (F) Representative IF images of RCAS-BRAF HGG with control (vehicle) and BRAFi+MEKi treatment and stained for GFAP, NESTIN, and DAPI (DNA); graph depicting quantification of marker-positive cells within the tumor field (right) **P<0.01 (unpaired t-test). n=4 mice per group. Mean±SEM. Scale bar: 120µm. (G) Representative IF images of RCAS-BRAF HGG with control (vehicle) and BRAFi+MEKi treatment and stained for MBP, PDGFRα, and DAPI (DNA); High-magnification images of the white-boxed areas are shown on the right; graph depicting quantification of marker-positive cells within the tumor field (right). *P<0.05 (unpaired t-test). n=4 mice per group. Mean±SEM. Scale bar: 170µm. (H) Schematic for the experimental timeline for assessing BRAFi+MEKi effects on BRAF^V600E^ HGG cell lines *in vitro*. (I-J) Representative ICC images of murine BRAF-M34 (I) and RCAS-BRAF (J) cell lines in spontaneous differentiation medium treated with vehicle or concomitant BRAFi+MEKi stained for MBP, OLIG2, NESTIN and DAPI for DNA. *P<0.05 by unpaired multiple t-tests and Holm- Šídák method. n=2 independent experiments, mean±SEM.

In GBM, NESTIN and PROM1/CD133 are markers associated with a stemness state that contributes to chemoradiation resistance and drives tumor recurrence.^68,69^ We therefore investigated the effects of BRAFi+MEKi treatment on NESTIN and CD133/PROM1 cells in mice. Double-positive cell states decreased, as indicated by FC and IF, consistent with downregulated *Nestin* transcript with BRAFi+MEKi **(Figures S3C-E)**. The abundance of CD133/PROM1+ single-positive cells, however, increased while their overall abundance remained unchanged **(Figure S3C-D)**. This suggests that BRAFi+MEKi treatment reduces NESTIN+ but not CD133/PROM1+ cell states, the latter appears to be intrinsically resistant to treatment, aligning with increased *CD133/PROM1* transcript level in patient’s recurrent tumors after treatment **(Figure 1D)**, and prior reports of CD133+ cells involved in BRAFi recurrence and resistance. ^43^

Lastly, we tested BRAFi+MEKi effects on tumor cell states *in vitro* by treating murine (BRAF-M34, RCAS-BRAF) and human (STN-49, aGBM5 ^70^) BRAF^V600E^-mutant HGG cell lines with BRAFi+MEKi for 13 days. STN-49, specifically generated for this study, was derived from patient 49 before treatment **(Figures 3H, S3H, STAR Methods)**. Tumor cells were subsequently analyzed by immunocytochemistry (ICC) for markers for OL-like (MBP), pan-OPC/OL-like (OLIG2), and NSPC/CSC-like states (NESTIN). BRAFi+MEKi reduced tumor cell viability in all cell lines, and increased MBP+ and OLIG2+ cell abundance **(Figures 3I-J, S3I-J)**, consistent with a treatment-induced boost in glial differentiation (AC- and OL-like) cell states observed *in vivo*.

NESTIN+ cells, abundant at baseline (vehicle treatment) in all four cell lines, were unchanged with 13-day BRAFi+MEKi treatment in murine HGG cells, which is further corroborated by findings from the RCAS-BRAF model **(Figure 3F)**, and consistent with data in patients **(Figures 1D and 1F)**. These data indicated that cells with NSPC/CSC-like states initially decreased with treatment **(Figures 3B, 3D and S3C-E)**, were reinstated with treatment over time **(Figures 3I-J, S3I-J)**. Notably, mRNA and protein levels were not concurrent in the RCAS-BRAF tumors, and model-specific differences are evident as Nestin+ cells are more abundant in untreated RCAS-BRAF tumors **(Figure 2I)** compared to BRAF-2341 and BRAF-M34 tumors **(Figures 2D-E, Table S2)**.

These findings demonstrate that BRAFi+MEKi treatment temporarily suppresses NSPCs and reduces cells with pre-OPC and OPC-like states, while consistently heightening transitions towards glial cell differentiation states reminiscent of myelin-forming OL and AC in BRAF^V600E^-mutant HGG.

### BRAFi+MEKi Treatment Boosts Interferon Response Pathway Gene Expression Signature in BRAF^V600E^-mutant HGG cells

BRAF-altered glioma exhibit higher CD8^+^ T cell infiltration and MHC class I expression, when compared to BRAF wildtype glioma,^64,65^ suggestive of a robust anti-tumor immune response that could be exploited therapeutically in combination with BRAFi+MEKi. We therefore investigated whether BRAFi+MEKi alters immunomodulatory programs in BRAF^V600E^ -mutant HGG cells, using transcriptome analyses and qRT-PCR for validation. We first analyzed the whole transcriptomes of BRAF^V600E^-mutant patient-derived cell lines (STN-49, aGBM5) and murine HGG cells (BRAF-2341) for differential gene expression and performed Reactome pathway analysis. We observed that 48-hour BRAFi+MEKi treatment upregulated a 67 transcript-interferon gamma (IFN-γ) receptor-associated signature across human and mouse transcriptomes **(Figures 4A-B, S4A-C, and S4F)**. Transcripts for Major Histocompatibility Complex (MHC) genes were included in this signature, along with other inflammation-related transcripts such as those encoding the signal transducer and activator of transcription (STAT) proteins. qRT-PCR confirmed upregulation of selected MHC class I/II genes (*HLA-A*, *HLA-B*, *HLA-DRA*, and *CIITA*) in patient-derived and murine cell lines **(Figures S4D, S4E, and S4G, Table S3)**. These data suggested that BRAFi+MEKi treatment alters the immunomodulatory activity of HGG cells, heightening antigen presentation by MHC class I/II molecules, and potentially boosting anti-tumor immune responses.

**Figure 4.**
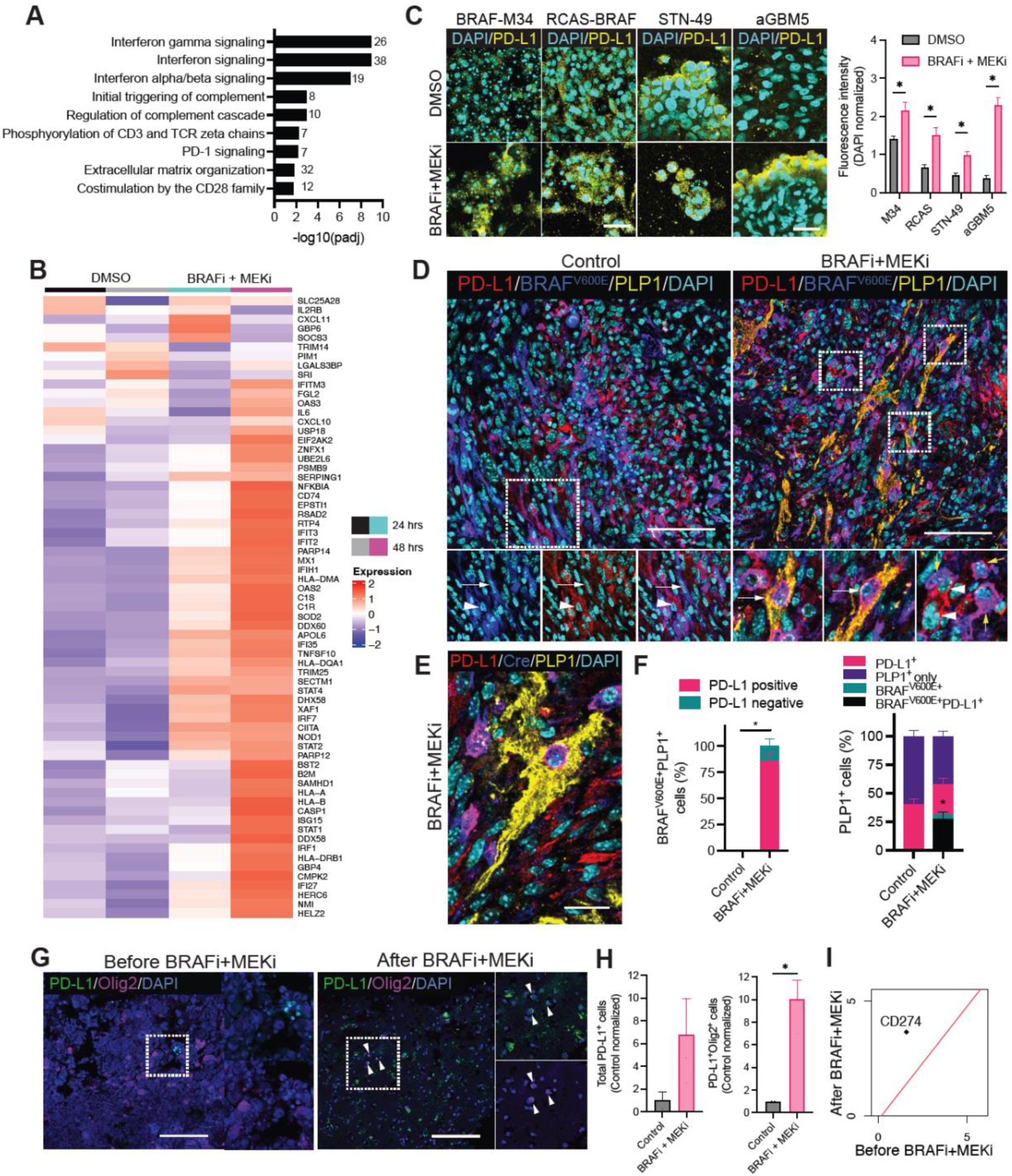
BRAFi+MEKi Treatment Induces Interferon Response Pathway Gene Expression Signature and Glial Differentiation Cell State Transitions, with a PD-L1 expressing immunomodulatory subpopulation. (A) Reactome pathway analysis revealed upregulated genes associated with interferon receptor signaling after 48-hour BRAFi+MEKi treatment to STN-49 cell line. Numbers on the bar graph indicate the gene count. (B) Heatmap demonstrating changes in expression of IFN-γ receptor-related transcripts after 24 and 48-hour BRAFi+MEKi treatment to STN-49 cell line. (C) Representative ICC images of murine and patient-derived tumor cells treated with BRAFi+MEKi and vehicle (DMSO) for 13 days. Quantification of (C). *P<0.05 (multiple paired t-tests). n=2 independent experiments. Mean±SEM. Scale bar: 40µm. **(D)** Representative IF images of BRAF-M34 HGG stained for PD-L1, BRAF V600E, PLP1, and DAPI (DNA). Control panel: BRAF^V600E^-mutated tumor cell negative for PD-L1 (arrow). PD-L1-positive non-tumor cell (arrowhead). BRAFi+MEKi panel: differentiated BRAF^V600E^-mutated glioma cells with PLP1 and PD-L1 co-expression (white arrows). PLP1-negative, PD-L1-positive glioma cells (yellow arrows). PD-L1 only positive cells (arrowheads). Scale bars: 70µm (left), 80µm (right). **(E)** Representative IF image of BRAFi+MEKi-treated BRAF-M34 HGG stained for PD-L1, Cre, PLP1, and DAPI. Differentiated Cre-positive glioma cell co-expressing PLP1 and PD-L1. Scale bar: 20µm. (F) Quantification of (D). Proportions of BRAF^V600E^ and PLP1 double-positive differentiated glioma cells with OL-like state positive or negative for PD-L1 (left panel), and proportions of PLP1-positive cells co-expressing BRAF^V600E^ and/or PD-L1 (right panel) were measured and compared between groups. *P<0.05 for BRAF^V600E+^PLP1^+^PD-L1^+^ triple-positive cells between treatment groups (two-way ANOVA with Sidak’s multiple comparisons test). n=3 mice per group. Mean±SEM. (G) Representative IF images of human BRAF^V600E^-mutant HGG (patient 49) tissue before and after BRAFi+MEKi treatment stained for PD-L1, Olig2, and DAPI (DNA). Scale bars = 180µm. (H) Quantification of (G). n=4 images from 2 consecutive sections of patient 49 tissue. Mean±SEM. (I) The scatter plot of *CD274* transcript (PD-L1) upregulated after BRAFi+MEKi treatment in patient 12.

### BRAFi+MEKi induces Glial Differentiation Cell State Transitions, with a PD-L1 expressing immunomodulatory subpopulation

IFN-γ signaling induces the PD-1 (Programmed Cell Death Protein-1) immune checkpoint pathway, which is frequently hijacked by cancer cells to circumvent anti-tumor immune responses and is an important immunotherapy target.^71^ We therefore conducted a comprehensive analysis of BRAFi+MEKi treatment effects on PD-1 signaling through Reactome and gene pathway analyses of whole transcriptomes, validated by ICC and IF. PD-1 signaling was one of pathways significantly altered by BRAFi+MEKi treatment, as shown by Reactome pathway analysis of human and murine BRAF^V600E^-mutant HGG cell line transcriptomes **(Figures 4A, S4H, S5A)**. PD-1 signaling components, including T-cell receptor complex protein *CD247,* and the PD-1 ligand *PD-L1* (*CD274*), were upregulated, as shown by gene network analysis **(Figure S5B)**. Next, we rigorously tested whether PD-L1 protein and gene expression aligned, using ICC and IF. BRAFi+MEKi treatment indeed increased PD-L1 protein expression in BRAF^V600E^-mutant HGG cells *in vitro* **(Figure 4C)**, and *in vivo* **(Figure 4D-F)**, consistent with upregulation of *PD-L1* transcript. We also found upregulated PD-L1 expression at protein and transcriptomic levels in BRAF^V600E^ HGG patients post-BRAFi+MEKi treatment **(Figures 4G-I)**. PD-L1 is also expressed in murine BRAF-2341 HGG, accounting for 83% of NESTIN+ cells **(Figure S4I)**.

Immunomodulatory roles for disease-associated OL-lineage cells have been reported, aside from their function in myelination.^72–75^ We then investigated if OL-like cell states enriched in response to BRAFi+MEKi treatment express the immunomodulatory PD-L1, by assessing the co-expression of PD-L1 with OL markers PLP1, MBP, and OLIG2, by IF and ICC. BRAFi+MEKi treatment significantly upregulated PLP1 expression in BRAF-M34 glioma (BRAF^V600E+^PLP1^+^) after 12 days of treatment **(Figures 4D-F)**, consistent with a boost in OL-like cell states **(Figure 3I)**. The vast majority (85.92%) of these OL-like tumor cells expressed PD-L1 **(Figure 4F, left panel)**, suggesting that the treatment induced immunomodulatory function in OL-like HGG cells. Interestingly, 40.28% of PLP1^+^ BRAF^V600^-negative cells were immunoreactive for PD-L1 even before treatment (vs. 25.87% after treatment), indicating these non-mutated OLs, also called glioma-associated OLs, might be hijacked by glioma cells to promote immunosuppression. Co-expression of MBP/Olig2 and PD-L1 was assessed *in vitro*, to complement our *in vivo* results. Positive fold changes for co-expression in murine and human cell lines are variable, suggesting model-specific differences **(Figure S4K, left panel)**. Lastly, co-expression of pan-OL marker OLIG2 and PD-L1 in BRAF^V600E^ HGG patient tumors revealed a 10-fold increase in OLIG2/PD-L1 double-positive cells after BRAFi+MEKi treatment **(Figure 4G-H)**.

In conclusion, our data provide compelling evidence that BRAFi+MEKi treatment induced a transition to differentiating glia cell (AC-like and OL-like) states, with a subpopulation of OL-like glioma cells acquiring immunomodulatory properties via PD-L1 expression, establishing a direct mechanistic link of cell state transitions and immune evasion.

### PD-L1 Expression Is Elevated in BRAF-mutant HGG and GBM Cells

Our data demonstrated that BRAFi+MEKi upregulated PD-L1 expression, a mechanism for T cell suppression and a key criterion for patient enrollment in PD-1 inhibition therapy.^76^ Therefore, we evaluated PD-L1 expression in a large cohort of patients with BRAF-mutant glioma, including those with BRAF^V600E^. First, we evaluated the association between PD-L1 expression and BRAF mutation status in a large cohort of adult GBM (IDH-wildtype) specimens (n=3126), undergoing different characterization analyses, including whole transcriptome sequencing and PD-L1 IHC analysis **(STAR Methods)**. We found that compared to BRAF-wildtype GBM patients (n=3043), those with BRAF-mutant/fused tumors (n=59) were younger (54 vs. 59 years, P=3.5x10^-3^), and likely to be MGMT-unmethylated (75% vs. 56%, P=2.1x10^10^). PD-L1 expression by IHC was significantly higher in BRAF-mutant GBMs (54%) than BRAF-wildtype (17%, P<0.0001) **(Figure 5A)**. This was orthogonally validated using whole-transcriptome sequencing data, showing increased PD-L1 (*CD274)* gene expression in BRAF-mutated (n=17) versus BRAF-wildtype (n=1022) GBMs (P=0.045) **(Figure 5B)**. We found that PD-L1 and BRAF V600E showed a high degree of co-expression **(Figure 5C)**, using immunohistopathology, further corroborating PD-L1 expression in BRAF^V600E^-mutant HGG cells. We also showed that PD-L1 is expressed in various BRAF^V600E^-mutant glioma types, including ganglioglioma, pleomorphic xanthoastrocytoma (PXA), and malignant astrocytoma/glioma, albeit with a varied frequency score, as determined by IHC **(Figure S5C, Table S4)**

**Figure 5.**
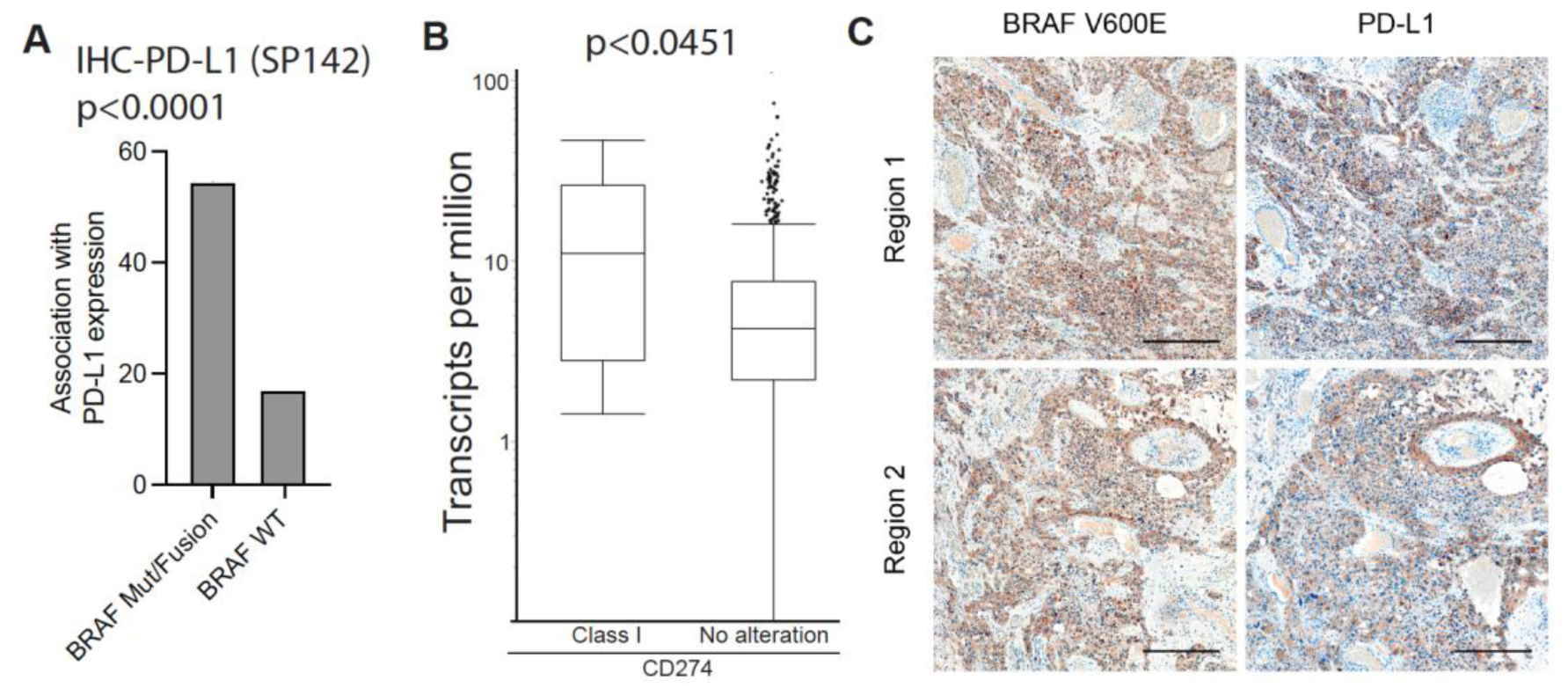
PD-L1 Expression is Elevated in BRAF-mutant HGG and GBM Cells. (A) Bar graph depicting percentage of patients with PD-L1+ tumors in the BRAF mutation/fusion versus BRAF wildtype (no alterations) groups (IHC). n=59 patients BRAF mut/fus and 3126 total patients (chi-squared tests). (B) Box plot representing *CD274* (PD-L1) transcripts per million in the BRAF mutation/fusion (n=17) versus BRAF wildtype (no alterations; n=1022) groups, from whole transcriptome sequencing (Wilcoxon Rank-Sum test). (C) Representative IHC images for BRAF V600E and PD-L1 in consecutive sections of BRAFV600E-mutant HGG. Scale bars: 250µm.

Our findings demonstrate that BRAF mutations, including the BRAF^V600E^ variant, are associated with elevated PD-L1 expression in GBM and other glioma subtypes. This correlation suggests that patients with BRAF-mutant gliomas may harbor suppressed T cells that are reactivatable through anti-PD-1 therapy, potentially eliciting a robust anti-tumor immune response and driving a significant survival benefit.

### BRAFi+MEKi Treatment Enhances Tumor-infiltrating T Cells

The efficacy of T cell checkpoint therapy is significantly influenced by the presence of tumor-infiltrating T cells. ^77^ ^78^ Since BRAF-altered glioma exhibits robust CD8^+^ T cell infiltration,^64,65^ and our data showed that they express PD-L1 **(Figures 4A-B, 4F, and 4H)**, we closely assessed tumor-infiltrating T cells in BRAF^V600E^-mutant murine glioma with and without BRAFi+MEKi treatment, using high-dimensional single-cell mass cytometry (CyTOF), scRNA-seq, and validation by IF **(Figure 6A, STAR Methods)**. We identified robust infiltration of CD4+ and CD8+ effector T cells (T_eff_), and CD4+FoxP3+ regulatory T cells (T_reg_) within the tumor mass, as confirmed by CyTOF (**Figures 6B, S6A-D),** scRNA-seq **(Figures 6G-H)** and IF **(Figures 6I, 6J, 6M, S5D, S6G)**. The presence of tumor-infiltrating T cells in murine BRAF^V600E^-mutant HGG is consistent with a heightened frequency of CD3+CD8+ T cells in human BRAF-altered glioma compared to BRAF wildtype astrocytoma.^79,80^ BRAFi+MEKi treatment further elevated the frequency of CD4+ and CD8+ T cells in all three models **(Figures 6B, 6C, 6H-J, 6M, S6G)**, consistent with upregulation of the antigen presentation machinery **(Figures 4B and S4B-G)** and suggestive of anti-tumor immunity heightened by treatment.

**Figure 6.**
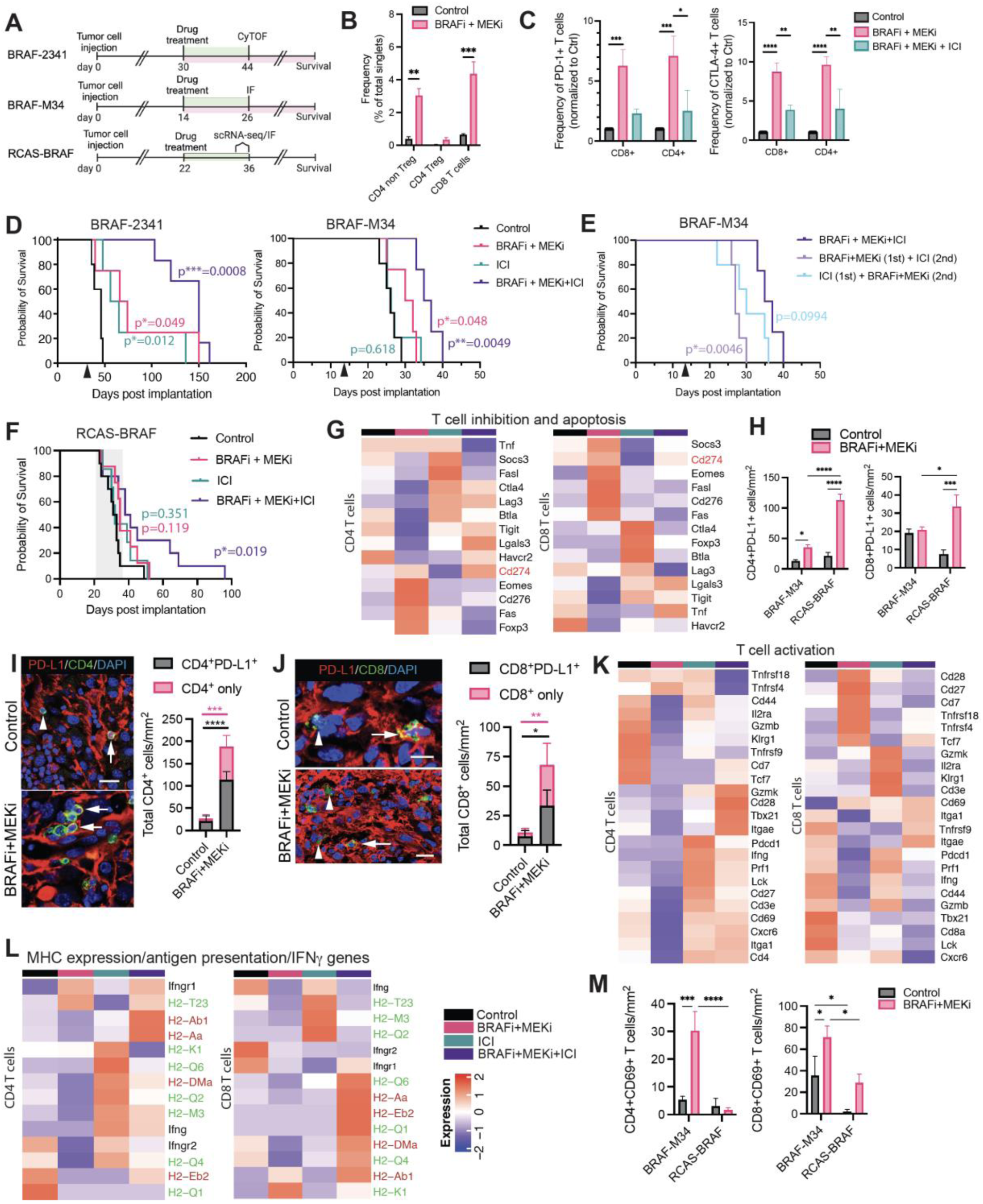
BRAFi+MEKi Enhances T Cell Infiltration and Synergizes with ICI to Improve T Cell-Dependent Survival. **(A)** Experimental timeline (green: 12∼14 days treatment for CyTOF, IF, and scRNA-seq analyses; pink: continuous treatment for survival analysis) in all three murine models. (B) Frequency of populations of tumor-infiltrating lymphocytes in BRAF-2341 tumors by CyTOF analysis. 3 mice/group and two independent experiments were analyzed. **P<0.01, ***P<0.001 (2-way ANOVA with Sidak’s multiple comparisons test). **(C)** Graphs depicting the frequency of PD-1+CD8+ and PD-1+CD4+ (left) or CTLA-4+CD8+ and CTLA-4+CD4+ (right) T cells normalized to control by CyTOF analysis. Data representative of two independent experiments and 3 mice/group. *P<0.05, **P<0.01, ***P<0.001, ****P<0.0001 (2-way ANOVA with Tukey’s multiple comparisons test). **(D)** Kaplan-Meier survival curve of syngeneic, orthotopic BRAF-2341 (left) or BRAF-M34 (right) HGG-bearing mice of all four treatment groups: 1) control (vehicle), 2) BRAFi+MEKi, 3) ICI, and 4) BRAFi+MEKi+ICI. Treatment initiation at day 30 (BRAF-2341) or day 14 (BRAF-M34) post-injection (black arrow). Data representative of two independent experiments and 5-6 mice/group (BRAF-2341) and 4-5 mice/group (BRAF-M34). P values are indicated in the graph and Tables 1-2 when compared to the control group. (E) Kaplan-Meier survival curve in immunocompetent C57Bl/6 mice with orthotopic BRAF-M34 treated with BRAFi+MEKi and ICI, either concurrently (BRAFi+MEKi+ICI) or sequentially. Treatment initiation at day 14 post-injection (black arrow). 4-5 mice/group. P values are indicated in the graph and Table 3 when compared to the concurrent treatment group. **(F)** Kaplan-Meier survival curve in immunocompetent C57Bl/6 mice with orthotopic RCAS-BRAF HGG treated with BRAFi+MEKi and/or ICI for 14 days. Treatment period from day 22 to day 36 post-injection (shaded area). Data representative of two independent experiments and 7-10 mice/group. P values are indicated in the graph and Table 1 when compared to the control group. (G) Heatmaps of T cell inhibition- and apoptosis-associated transcripts in CD4+ and CD8+ T cells in RCAS-BRAF model. (H) Densities of CD4+PD-L1+ and CD8+PD-L1+ T cells in BRAF-M34 and RCAS-BRAF (IF analysis). *P < 0.05, ***P<0.001, ****P<0.0001 (two-way ANOVA with uncorrected Fisher’s LSD). n=4 mice per group. Mean±SEM. **(I-J)** Representative IF images of RCAS-BRAF tissue stained for PD-L1, CD4 (I), and CD8 (J). CD4+ and CD8+ cells positive for PD-L1 (arrows); CD4+ and CD8+ cells negative for PD-L1 (arrowheads). Quantifications of cell populations after BRAFi+MEKi treatment compared with vehicle control treatment. *P < 0.05, **P<0.01, ***P<0.001, ****P<0.0001 (two-way ANOVA with Sidak’s multiple comparisons test). n=4 mice per group. Mean±SEM. Scale bars: (I) 20µm (top), 15µm (bottom), (J) 10µm (top), 20µm (bottom). (K) Heatmaps of T cell activation-associated transcripts in CD4+ and CD8+ T cells in the RCAS-BRAF model. (L) Heatmaps of MHC class I (green), II (red) and IFNγ (black)-related transcripts associated with antigen presentation in CD4+ and CD8+ T cells in the RCAS-BRAF model. **(M)** Densities of CD4+CD69+ and CD8+CD69+ T cells from BRAF-M34 and RCAS-BRAF (IF analysis). *P < 0.05, ***P<0.001, ****P<0.0001 (two-way ANOVA with uncorrected Fisher’s LSD). n=3-4 mice per group. Mean±SEM.

**Table 1.** Number of mice, medial survival, and statistical analyses by the log-rank test in Fig 5D (FVB/N:BRAF-2341), Fig S8E (nu/nu:BRAF-2341) and Fig 5F (C57BL/6:RCAS-BRAF).

| Host Strain | FVB/N |  |  | nu/nu |  |  | C57BL/6 |  |  |
| --- | --- | --- | --- | --- | --- | --- | --- | --- | --- |
| Cell line | BRAF-2341 cell line |  |  | BRAF-2341 cell line |  |  | RCAS-BRAF cell line |  |  |
| Group | n | Median Survival | P value | n | Median Survival | P value | n | Median Survival | P value |
| Control | 5 | 46 | — | 5 | 19 | — | 10 | 32 | — |
| BRAFi+MEKi | 5 | 70 | *0.049 | 5 | 30 | *0.025 | 8 | 35 | 0.119 |
| ICI | 5 | 60.5 | *0.012 | 6 | 21.5 | 0.189 | 7 | 32 | 0.351 |
| BRAFi+MEKi+ICI | 6 | 150 | ***0.0008 | 6 | 32.5 | *0.01 | 10 | 39.5 | *0.019 |

**Table 2.** Number of mice, medial survival, and statistical analyses by the log-rank test in Fig 5D (non-T cell-depleted C57BL/6: BRAF-M34) and Fig S8E (T cell-depleted C57BL/6: BRAF-M34).

| Host Strain | C57BL/6 |  |  | C57BL/6<br>with anti-CD4 and anti-CD8 Abs |  |  |
| --- | --- | --- | --- | --- | --- | --- |
| Cell line | BRAF-M34 cell line |  |  | BRAF-M34 cell line |  |  |
| Group | n | Median Survival | P value | n | Median Survival | P value |
| Control | 5 | 26 | — | 5 | 21 | — |
| BRAFi+<br>MEKi | 5 | 31 | *0.0485 | 4 | 28 | *0.0018 |
| ICI | 4 | 26 | 0.6182 | 5 | 21.5 | 0.4333 |
| BRAFi+<br>MEKi+<br>ICI | 4 | 36 | **0.0049<br>vs. control<br>*0.0169<br>vs.<br>BRAFi+<br>MEKi | 4 | 30 | *0.0046<br>vs. control<br>0.5296<br>vs.<br>BRAFi+<br>MEKi |

### Combination of BRAFi+MEKi with ICI Enhances Survival Outcomes in a T Cell-Dependent Manner

Our findings provided a strong rationale for combining BRAFi+MEKi with PD-1/PD-L1 immune checkpoint inhibition (ICI) to boost T-cell activation and improve survival. To explore this, we assessed the effects of BRAFi+MEKi combined with ICIs on the T cell compartment in our three mouse models **(Figure 6A)**. Mice bearing orthotopic BRAF^V600E^-mutant HGG tumors were treated with dabrafenib and trametinib (BRAFi+MEKi), and/or anti-PD-L1 antibody. Since PD-1/PD-L1 inhibitors alone have shown no survival benefit in HGGs,^57,58^ we also included CTLA-4 blockade (α-PD-L1+α-CTLA-4; referred to as ICI) **(Figure 6A and S5E)**. Concurrent, quadruple treatment (BRAFi+MEKi+ICI) significantly improved survival over BRAFi+MEKi or ICI alone in all three models **(Figures 6D-F**, **Tables 1-2).** This survival benefit is associated with T cell reactivation, as shown by reduced frequencies of PD-1– and CTLA4–expressing CD4+/CD8+ T cells **(Figure 6C)** and T_reg_ cells **(Figures S6A-D)**.

To assess model-specific differences for BRAF-M34 and RCAS-BRAF further, we analyzed the frequency of CD4+ and CD8+ T cells co-expressing inhibitory (PD-L1/PD-1) and activation (CD69) markers. BRAFi+MEKi treatment increased PD-L1+ and PD-1+ T cell frequencies **(Figure 6H-J, S6F, S6H)**, suggesting that the PD-L1/PD-1 checkpoints become activated to escape BRAFi+MEKi-induced T cell activation. Only PD-1+ T cells in the RCAS-BRAF model showed the opposite trend, most likely undergoing apoptosis after treatment **(Figures 6G, S6H)**.

We observed differences in T cell inactivity across the models. RCAS-BRAF showed the highest T cell suppression, while BRAF-2341 had the least (**Figure S6F**). This variability is expected and consistent with reported model-specific differences and differences in the immune responses even in genetically identical mice **(Figures 6D-F)**. Interestingly, both BRAF-M34 and RCAS-BRAF models did not respond to ICI treatment alone, suggesting their T cells were unresponsive to activation **(Figures 6D, 6F, 6H).** However, their T cells were responsive to BRAFi+MEKi alone, with the BRAF-M34 model showing significantly higher CD4+CD69+ and CD8+CD69+ T cells, indicative of early T cell activation, compared to the RCAS-BRAF model **(Figure 6M).** Furthermore, our *in vitro* data showed that BRAF-M34 glioma cells with OL-like states have less immunosuppressive effect on T cells **(Figure S4K)**.

The survival benefits seen with therapeutic antibodies, whether alone or in combination with BRAFi+MEKi, were completely abolished in athymic mice lacking T cells. Similar results were observed in BRAF-M34 mice that are depleted of T cells, confirming T cell-dependent anti-tumor efficacy **(Figure S6E, Tables 1-2)**. Although T cells are not strictly required for the anti-tumor effects of BRAFi+MEKi, increased T cell activation and anti-tumor immunity may contribute to the treatment’s effectiveness, particularly in the BRAF-M34 model.

Lastly, we tested whether sequential therapy optimizes BRAFi+MEKi responses, similar to what has been reported for BRAF^V600E^-mutant melanoma patients.^81^ ICI 1^st^ showed a trend for improved survival over BRAFi+MEKi 1^st^, which provided no survival benefit over BRAFi+MEKi treatment alone. Importantly, concurrent quadruple treatment provided the greatest survival benefit and is superior to any sequential approach **(Figures 6D-E**, **Table 3)**. These findings suggest that patients with BRAF^V600E^-mutant HGG are most likely to benefit from a combination treatment regimen integrating BRAFi+MEKi and ICIs concurrently.

**Table 3.** Number of mice, medial survival, and statistical analyses by the log-rank test in Fig 5E (C57BL/6: BRAF-M34, sequential treatment).

| Host Strain | C57BL/6 |  |  |
| --- | --- | --- | --- |
| Cell line | BRAF-M34 cell line |  |  |
| Group | n | Median Survival | P value |
| BRAFi+MEKi (first) + ICI (second) | 5 | 27 | *0.0046 vs. concurrent treatment<br>0.1508 vs. ICI (first) + BRAFi+MEKi (second) |
| ICI (first) + BRAFi+MEKi (second) | 5 | 30 | 0.0994 vs. concurrent treatment |

### Synergistic Effect of BRAFi+MEKi with ICI On T Cell Reactivation

Our data demonstrated that quadruple (BRAFi+MEKi+ICI) treatment overcomes resistance observed with dual therapies (BRAFi+MEKi and ICI alone) in the RCAS-BRAF model (Figure 6F). To gain insights into potential differences in T cell dynamics with dual and quadruple treatments, we analyzed CD4+/CD8+ T cell and tumor clusters in scRNA-seq data from the RCAS-BRAF model when mice reached their disease-related endpoint **(Figure 6A)**. BRAFi+MEKi moderately upregulated selected suppression markers on T cells, *including Cd274* (PD-L1), and robustly upregulated others, such as *Cd276, Eomes, Socs*, *Fas* and *Fasl* in both CD4+ and CD8+ T cell clusters **(Figure 6G)**. *Foxp3* was upregulated in CD4^+^ T cells post-treatment, indicating the expansion of T_reg_ cells. BRAFi+MEKi downregulated activation markers in CD4+ T cells, including *Gzmk*, *Gzmb*, *IL2ra*, and others, while upregulating immunosuppressive markers, including *Cd27*, *Cd7, Tnfrsf18* and *Tnfrsf4* in CD8+ T cells **(Figure 6K)**, suggesting the presence of dysfunctional CD8+ T cells after BRAFi+MEKi treatment.^82^

We also found that treatment reduced expression of MHC class I and II genes in CD4+ and CD8+ T cells **(Figure 6L)**. MHC class I gene expression was diminished in tumor cells and tumor-associated macrophages (TAMs)/microglia **(Figures S6J-K)**. This downregulation was suggestive of suppressed antigen presentation, potentially enabling immune evasion and BRAFi+MEKi-resistance at the survival endpoint.

ICI treatment alone upregulated T_reg_ expansion markers, including *Foxp3* and *Btla* **(Figures 6G and 6K)**. Moreover, increased expression of inhibitory markers like *Ctla4, Lag3,* and *Pdcd1* **(Figures 6G and 6K)** in CD4+ and CD8+ T cells, and increased MHC and *Ifng* expression, indicated activation balanced with suppression of T cell activity **(Figures 6K-L)**. ICI also caused excessive pro-inflammatory cytokine production in CD4+ T cells and tumor cells, resembling a ‘cytokine storm’ **(Figure S6I)**.

In contrast, quadruple treatment induced expression of genes linked to T cell activation and effector functions, including *Itgae, Gzmk, Cd28, Tbx21, Itga1, Cd69,* and *Tnfrsf9* **(Figure 6K).** Importantly, MHC class I/II gene expression was restored for effective antigen presentation without causing excessive cytokine production **(Figures 6L and S6I)**. MHC class II gene upregulation in CD8+ T cells is associated with memory T cell differentiation **(Figures 6L and S6D)**.^83^ This may contribute to a more durable anti-tumor immune response.

Our findings indicate that BRAFi+MEKi treatment resistance include T_eff_ cell suppression, T_reg_ expansion, and impaired antigen presentation, which collectively could lead to immune evasion in the RCAS-BRAF model. While ICI activated T cells, it also induced severe cytokine release syndrome (cytokine storm) due to excessive cytokine production. In contrast, quadruple treatment effectively ensures full activation of CD4+ and CD8+ T cells, which could be pivotal for overcoming therapy resistance and conferring a significant survival benefit.

### BRAFi+MEKi-Induced Galectin-3 Secretion Links Glial Differentiation To PD-L1 Upregulation

To investigate the mechanism for cell state transitions and PD-L1 expression by BRAFi+MEKi, we assessed the secretome of BRAF^V600E^-mutant HGG cells *in vitro* using a high-throughput nELISA assay. STN-49 and aGBM5 cell lines were treated with BRAFi+MEKi over 13 days, mimicking the duration of *in vivo* treatments **(Figure 7A, STAR Methods)**. Molecular factors measured by nELISA that were either upregulated or below the detectable threshold post-treatment are presented in a heatmap **(Figure S7A).** BRAFi+MEKi significantly elevates the secretion of the carbohydrate-binding lectin family protein Galectin-3^84^ **(Figure 7B)**, with levels peaking at day 13, coincidently with increased glial differentiation **(Figure S3I-J)**. RNA-seq confirmed upregulation of the Galectin-3 encoding gene *LGALS3* with 48 hour-treatment of BRAFi+MEKi **(Figures 7C-D)**. In addition to Galectin-3, other factors IL-23 and IL-32 were also upregulated post-treatment, albeit the timing of upregulation varied between STN-49 and aGBM5 cell lines **(Figure S7B)**. This indicates that cytotoxic response rates to BRAFi+MEKi treatment could be cell line-specific.

**Figure 7.**
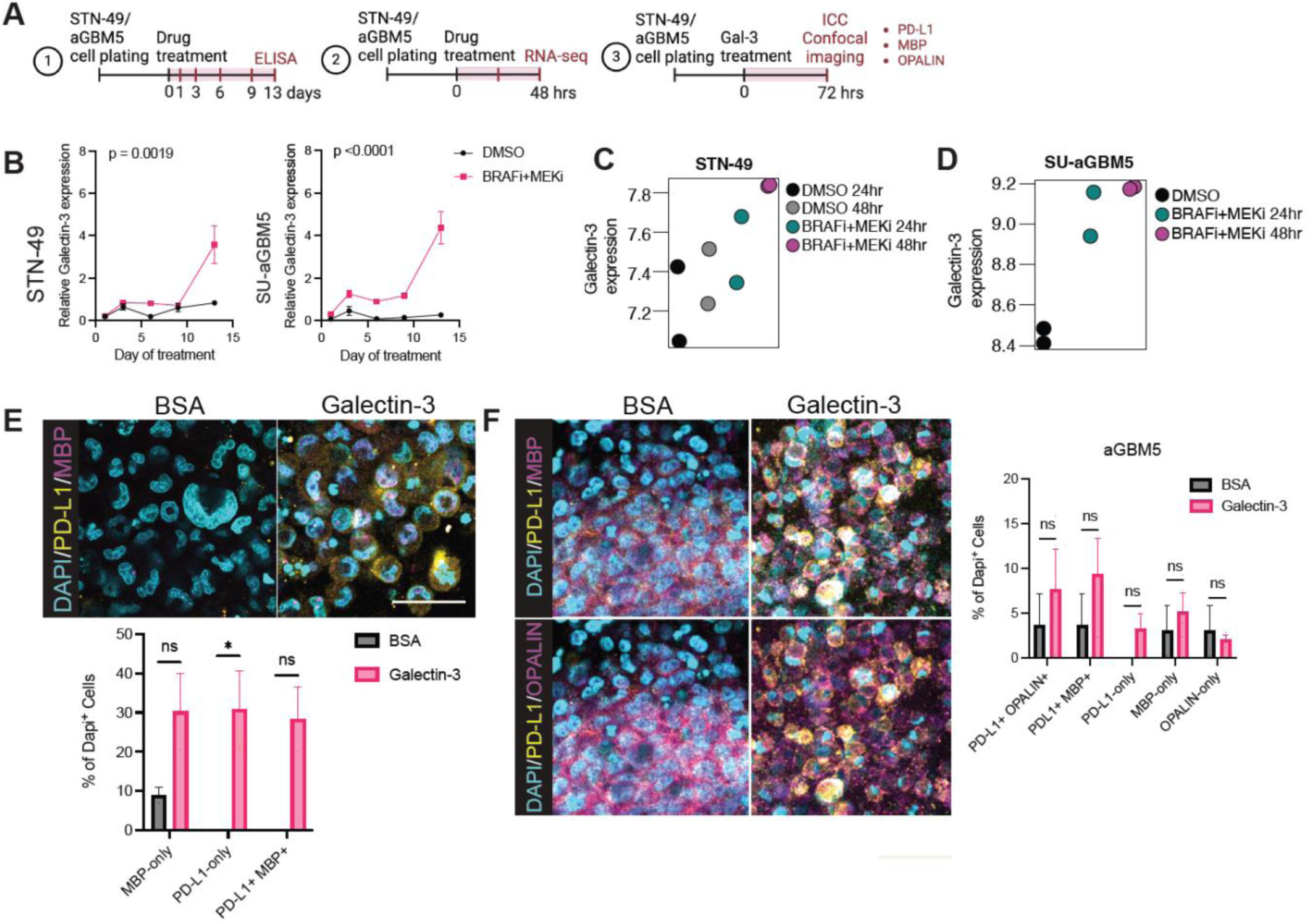
BRAFi+MEKi-Induced Galectin-3 Secretion Links Glial Differentiation To PD-L1 Upregulation. (A) Timelines of *in vitro* experiments for nELISA assay, bulk RNA-seq, and ICC using STN-49 and aGBM5 cell lines. (B) Relative Galectin-3 concentrations measured over time with nELISA in supernatants by STN-49 or SU-aGBM5 cell lines treated with BRAFi+MEKi or DMSO (two-way ANOVA with multiple comparisons). Simple linear regression ± SEM. (C-D) Scatter plots representing Galectin-3 expression levels in STN-49 (C) and aGBM5 (D) cell lines treated with DMSO-Control or BRAFi+MEKi at 24 and 48 hours. (E-F) Representative ICC images of STN-49 (E) and aGBM5 (F) cell lines treated with BSA or recombinant Galectin-3 protein for 72 hours and subsequently stained for PD-L1, MBP, or OPALIN and DAPI. Bar graphs showing cellular quantification (% of total DAPI+ cells). *P<0.05 (2-way ANOVA with Sidak’s multiple comparisons test). ns, not significant. n=2-3 replicates from one independent experiment. Mean±SEM. Scale bars: 40µm.

Next, we investigated a potential direct role for Galectin-3 in inducing glial differentiation cell state transitions and PD-L1 expression in STN-49 and aGBM5 cell lines **(Figure 7A)**. Adding recombinant Galectin-3 to cells in vitro efficiently boosted both glial differentiation cell states and PD-L1 expression in STN-49 cells **(Figure 7E),** whereas aGBM5 exhibited only an upward trend in PD-L1 expression in differentiated glial populations **(Figure 7F).** These findings suggest that Galectin-3 secretion from HGG cells potentially provides a key link of tumor differentiation and PD-L1 upregulation in response to BRAFi+MEKi treatment.

## DISCUSSION

BRAF and MEK inhibitors have revolutionized treatment for BRAF^V600E^ –mutant cancers, including gliomas, where molecular targeted therapies are otherwise limited. Despite these successes, treatment challenges persist, particularly in pLGG patients who experience tumor rebound after treatment cessation, and in HGG patients who develop therapy resistance.^85^ Resistance mechanisms across tumor types can be categorized as primary (intrinsic), adaptive, and acquired. While intrinsic and acquired resistance mechanisms to BRAFi and BRAFi+MEKi are well described in several solid cancers, like melanoma, the specific mechanisms underlying resistance in glioma remain poorly understood. ^32,86^ Our study fills a critical knowledge gap by investigating glioma lineage-specific resistance mechanisms. The intrinsic resistance to BRAFi+MEKi is higher for BRAF^V600E^-mutant HGG than melanoma, underscoring the importance of cancer lineage-specific studies. The high rates for intrinsic resistance of BRAF^V600E^-mutant HGGs may partially be explained by their higher intra-tumoral heterogeneity when compared with LGG. We showed earlier that a polo-like-kinase 1 (PLK1)-high subpopulation, characterized by prolonged G2-M phase, increased asymmetric cell division, and resistance to BRAFi, drives recurrence.^43,67^ Our data here convincingly corroborated that BRAF^V600E^-mutant HGGs express gene modules for neuronal and glial cells representing diverse differentiation states.

We found that BRAFi+MEKi treatment induces significant shifts in glioma cell state, favoring more differentiated phenotypes at the expense of stem and progenitor-like states. Our *in vitro* data further support this observation, showing that surviving cells adapt to treatment by acquiring a more differentiated state,^87^ which is associated with upregulated immunomodulatory PD-L1 expression. This phenotypic heterogeneity and cell plasticity may underlie the high intrinsic resistance and rapid adaptation to treatment observed in BRAF^V600E^-mutant HGGs. Indeed, phenotypic heterogeneity within tumor cells driven by tumor cell plasticity is emerging as a key feature of IHD-wildtype GBM, enabling GBM cells to transition between four major cell states. ^63^

Lineage-specific responses to BRAFi+MEKi treatment were also observed. CD133/PROM1, a marker for both CSCs and differentiating OLs,^62,88^ remains stable or increased with BRAFi+MEKi; whereas NESTIN+ cells decrease after short-term BRAFi+MEKi but persist with prolonged exposure. This may reflect the presence of distinct CSC populations with unclear roles in tumor recurrence. Notably, differentiated glioma cells have been shown to accelerate CSC-driven tumor growth,^89^ raising the possibility that BRAFi+MEKi-induced transition to glial cell states drives tumor recurrence by maintaining a CSC population. Importantly, therapy resistance is traditionally associated with cell de-differentiation, with stem cell-like states being related to therapy resistance in glioma. We provide comprehensive evidence that BRAFi+MEKi induces differentiation rather than de-differentiation, and suggest this transition to a more differentiated, immunosuppressive state may be a unique adaptation and resistance mechanism in glioma.

Moreover, acquired resistance mechanisms in glioma share some similarities with those in melanoma but also exhibit distinct lineage-specific differences. Analysis of paired pre-/post-treatment samples from BRAFi or BRAFi+MEKi-treated patients identified 13 putative resistance-associated alterations, including mutations in *ERRF1*, *BAP1*, *ANKHD1*, and *MAP2K1*. ^32^Despite a small patient cohort size (4 out of 15 with BRAFi+MEKi), only about 50% of cases show a *de novo* mutation, suggesting that non-genetic adaptive resistance mechanisms, such as tumor cell plasticity and cell state transitions described in our studies, may play an important role in resistance in BRAF^V600E^-mutant glioma. Indeed, our data here convincingly showed that BRAFi+MEKi induces cellular plasticity, which may serve as a therapeutic vulnerability. Further studies are needed to determine whether the adaptive mechanisms we uncovered also apply to non-BRAF^V600E^-mutant glioma and BRAF-wildtype glioma.

Our two novel murine models (BRAF-M34, RCAS-BRAF) recapitulate features of human BRAF^V600E^-mutant HGG, ^66^ including intra-tumoral cellular heterogeneity, with model-specific differences. These models provide robust preclinical tools, facilitating immunological studies due to their intact immune system and cross-model comparisons owing to their identical C57BL/6 genetic backgrounds.

Our data reveal that BRAFi+MEKi may induce adaptive resistance via increased Galectin-3 secretion from glioma cells, a response to stress conditions such as hypoxia,^90^ and, as implicated by our data – therapeutic insults. Tumor-secreted Galectin-3 governs several inter-connected processes, including MAPK reactivation ^90^, glial differentiation, and PD-L1 upregulation, all contributing to immune evasion. MAPK signaling plays multiple roles in glial cell differentiation^91^ and myelin maintenance.^92,93^ Through K-Ras/MEK signaling, Galectin-3 promotes cell survival, and proliferation in various cancers.^90^ Therefore, Galectin-3 could drive MAPK reactivation via K-Ras-Raf-ERK1/2 signaling,^90^ facilitating glioma cell differentiation while simultaneously upregulating PD-L1 as an adaptive response to BRAFi+MEKi. It is not well understood what drives cancer cell state transitions in glioma, although they are a well-recognized adaptative mechanism enabling immune evasion, therapy resistance, and metastasis in many cancer types,^94^ influencing both plasticity and the TME. We found that BRAFi+MEKi treatment induces Galectin-3 secretion, promoting glioma cell differentiation into AC- and OL-like states, mirroring its role in normal OL differentiation with increased MBP expression.^95,96^ Moreover, this differentiation is associated with increased PD-L1 expression, fostering an immunosuppressive TME. Given the antigen-presenting capabilities of ACs and OLs,^72,73,97,98^ these observed cell state transitions may actively shape immune responses in glioma, linking glioma differentiation and immune evasion.

Gliomas are considered immune-inactive due to their low mutational burden,^63^ but BRAF^V600E^-mutant gliomas are more immune-rich, with elevated CD8^+^ T cell infiltration, MHC class I expression^79^, and PD-L1 expression, suggesting intrinsic ICI sensitivity. Despite encouraging data in mouse models,^66,67^ clinical trials with PD-1 and/or CTLA-4 inhibitors show limited efficacy in GBM.^68,69^ Nevertheless, combined PD-1 blockade with MEKi shows promising immune synergism in a small number of patients.^57^ PD-1 and CTLA-4 blockade is also effective as second-line treatment after BRAFi+MEKi in BRAF^V600E^-mutant metastatic melanoma,^55,56^ providing a strong rationale for exploring BRAFi+MEKi+ICI combinations in glioma.

Our study demonstrates that concurrent quadruple therapy with BRAFi+MEKi and ICI significantly improves survival compared to BRAFi+MEKi or ICI alone across all preclinical mouse models, by preserving antigen presentation and enhancing T cell responses. In contrast to sequential therapy, concurrent quadruple therapy effectively activates CD4+ and CD8+ T cells without exacerbating immune-related toxicity, overcoming resistance and conferring a significant survival advantage. Combination therapy may also reduce toxicity due to opposing on-target, off-tumor effects of BRAF or MEK inhibition on MAPK signaling.

While clinical trials of BRAFi+MEKi+ICI in BRAF^V600E^ HGG are lacking, melanoma studies provide insights. Patients previously treated with BRAFi+MEKi showed similar outcomes when subsequently treated with PD-1 or PD-1+Ipilimumab, and the combination of PD-1+Ipilimumab had higher toxicity compared to PD-1 alone. ^55^ Subsequent BRAFi+MEKi treatment after anti-PD-1 monotherapy doubled ORR compared to anti-CTLA-4 treatment alone.^99^ This is consistent with our survival analysis of sequential treatment, which shows that sequential ICI followed by BRAFi+MEKi offers benefit, though less than concurrent quadruple treatment. Additionally, BRAF-wildtype patients responded better to anti-PD-1+MEKi, while BRAF-mutated patients had lower ORR due to acquired resistance to treatment.^100^

Toxicity remains a concern, as BRAFi+MEKi causes fever, leukopenia, and hyponatremia, while ICIs induce immune-related adverse effects. ^27,101^ The principal reason for stopping sequential treatment (ICI 1^st^, BRAFi+MEKi 2^nd^) was disease progression rather than adverse effects, which is promising and could support quadruple treatment. Notably, dose reductions for BRAFi+MEKi could be considered for patients on quadruple treatment when tumors respond.^56^ Quadruple treatment may increase toxicity, but pediatric patients may tolerate it better than adults, and dose reductions or sequential treatment combined with Galectin-3 inhibitors could help balance efficacy and safety. Clinical data in metastatic melanoma and glioma highlight the necessity for further preclinical studies to refine treatment timing and minimize toxicity in HGG.

Overall, our study underscores the necessity of implementing concurrent quadruple therapy to generate more durable responses, thereby mitigating resistance mechanisms in BRAF^V600E^-mutant HGG and GBM.

In our preclinical models, BRAFi+MEKi could synergize with ICI by increasing T cell activity and antigen presentation, thus enhancing the intrinsic sensitivity of BRAF^V600E^ HGG.^102^ We hypothesize that sequential treatment (BRAFi+MEKi 1^st^, ICI 2^nd^) is less effective because it allows time for tumor cells to suppress T cell activity to reduce ICI efficacy, potentially through the secretion of Galectin-3.^102^ **(See hypothetical model, Figure S9D)**. Future studies will address whether Galectin-3 inhibition could further improve therapeutic outcomes by simultaneously targeting adaptive resistance and immunosuppressive TME. In summary, our study underscores the necessity of implementing concurrent quadruple therapy to generate more durable responses, thereby mitigating resistance mechanisms in BRAF^V600E^-mutant HGG and GBM. Incorporating Galectin-3 inhibitors into existing treatment regimens for these gliomas also offers a promising strategy to enhance therapeutic efficacy while managing toxicity, thereby improving the overall quality of life for patients.

### Limitations of the study

The rarity of BRAF^V600E^-mutant HGG limited the availability of matched pre- and post-treatment patient samples. To compensate for this, we generated two patient-derived cell lines and three immunocompetent mouse models/murine cell lines that effectively recapitulate multiple aspects of the human disease.^103^ Variability in treatment response within models was noted, this inter-individual variability is expected in immunomodulatory treatments.^104–106^ Despite model-specific differences, concurrent quadruple treatment consistently improved survival across all models, although the RCAS-BRAF model showed limited response, which may be caused by a higher proportion of PD-L1 expressing T cells **(Figure S8F),** leading to an increased extent of drug resistance. Nevertheless, future studies are required to elucidate resistance mechanisms underlying the limited responsiveness in this model to optimize the efficacy of concurrent treatment. Another limitation is that we did not fully characterize BRAFi+MEKi-induced cytokine alterations in tumor cells and/or their effects on CD4/CD8 T cell activation, which should be explored in future studies. Lastly, variability in drug responses across different patient-derived cell lines may reflect intrinsic differences in tumor cell types or Galectin-3 receptor expression of the cell lines, suggesting the need for single-cell analyses and expanded biological replicates or patient-derived cell lines.

## STAR METHODS

Detailed methods are provided in the online version of this paper and include the following

Key Resources Table

**Resources Availability**

Lead contact Materials availability

Data and code availability

Experimental Model and Subject Details

Animal Studies Human Studies

Cell lines and Primary Cultures

Intracranial Injections of Adenovirus

Orthotopic Glioma Generation

Drug Preparation and Treatment T cell Depletion

Magnetic Resonance Imaging

Tissue Collection and Processing of Mouse Samples

Immunofluorescence Staining on Frozen Tissue Sections of Mouse and Human Samples

Hematoxylin & Eosin Staining, Immunohistochemistry and Bulk RNA Sequencing

Differentiation Assays

nELISA Assay Immunocytochemistry

Confocal imaging and image processing Flow Cytometry and MACS

Mass Cytometry and Analyses

cDNA Synthesis and Quantitative PCR RNA Sequencing and Analysis

Single-cell Library Preparation and Sequencing

Analysis of Single-cell RNA Sequencing Data

**Statistical Analysis**

List of Supplementary Materials:

Figures S1 – S7.

Tables S1 – S4.

## Acknowledgments

We would like to thank Sista Sugiarto, Anne Marie Barrette, Daniella Morales, Kevin Cordero, and Caitlynn Tran for their excellent technical assistance. We thank Egbert Lu, Bill Weiss, Erin Simmons, and the UCSF Flow Cytometry Core for sharing the mass cytometry panel and running the mass cytometry analyses and the UCSF Brain Tumor Tissue Core and Stanford Neuroscience Tissue Bank for providing patient samples. We also thank Mr. Steve Avolicino from Histo-Tec Laboratory Inc. for tissue processing and immunohistochemical staining. We acknowledge the use of Stanford facilities including Stanford Veterinary Service Center, Neurosciences Preclinical Imaging Lab (Dr. Jieun Kim), Neuroscience Microscopy Service (Dr. Gordon Wang), Human Pathology/Histology Service Center (Pauline Chu), Stanford Human Immune Monitoring Centre (Molly Miranda), and Anita Metha-Damani and Xue Gong (Synthekine) for advice on T cell profiles.

## Funding

This work was supported by the

National Institute of Health R21 NS099836 (C.K.P)

National Institute of Health R01 NS080619 (C.K.P)

National Institute of Health R01 CA164746 (C.K.P.)

National Institute of Health 17X074 (C.K.P.)

Neurosciences Preclinical Imaging Lab Pilot Grant 2022 (C.K.P.)

Stanford Women’s Health and Sex Differences Center (WHSDM)

Seed Grant (C.K.P)

BRAF LGG consortium research fund (C.K.P; K.M.W.)

Stanford Cancer Institute (C.K.P)

National Institute of Health U54 CA 261717 Shurl and Kay Curci Foundation (L.M.P.)

Stanford Maternal & Child Health Research Institute (L.M.P.)

Chambers-Okamura Faculty Scholar in Pediatric Neurosurgery (L.M.P).

National Institutes of Health P30AG066515 (J.J.N)

## Author Contributions

Conceptualization: CKP, SG

Data acquisition: YLX, DP, SG, JWP, BB, ZPF, RW, PC, PH, JW, JX, AB

Data Analysis: YLX, DP, SG, JWP, BB, JJN, ZPF, KK

Funding acquisition: CKP, LMP

Investigation: YLX, DP, BB, JJN

Methodology: YLX, DP, ZPF, AB

Project administration: CKP

Resources: CKP, DH, KMW, JMML, KF, PNH, XJ, LMP, GAG, ML

Supervision: CKP

Writing-original draft: CKP, YLX, DP, JWP

Writing-review & editing: CKP, YLX, DP

## Competing Interests

Authors declare that they have no competing interests.

## Notes

### Competing Interest Statement

The authors have declared no competing interest.

### Summary of Updates

During the revision, we uncovered new findings linking cell state transitions to PD-L1 upregulation, as a mechanism for immune evasion which could potentially be mediated by galactin-3 secretion from glioma cells.

