## Supplemental Information for "BRAF/MEK Inhibition Induces Cell State Transitions Boosting Immune Checkpoint Sensitivity in BRAF^V600E^-mutant Glioma"

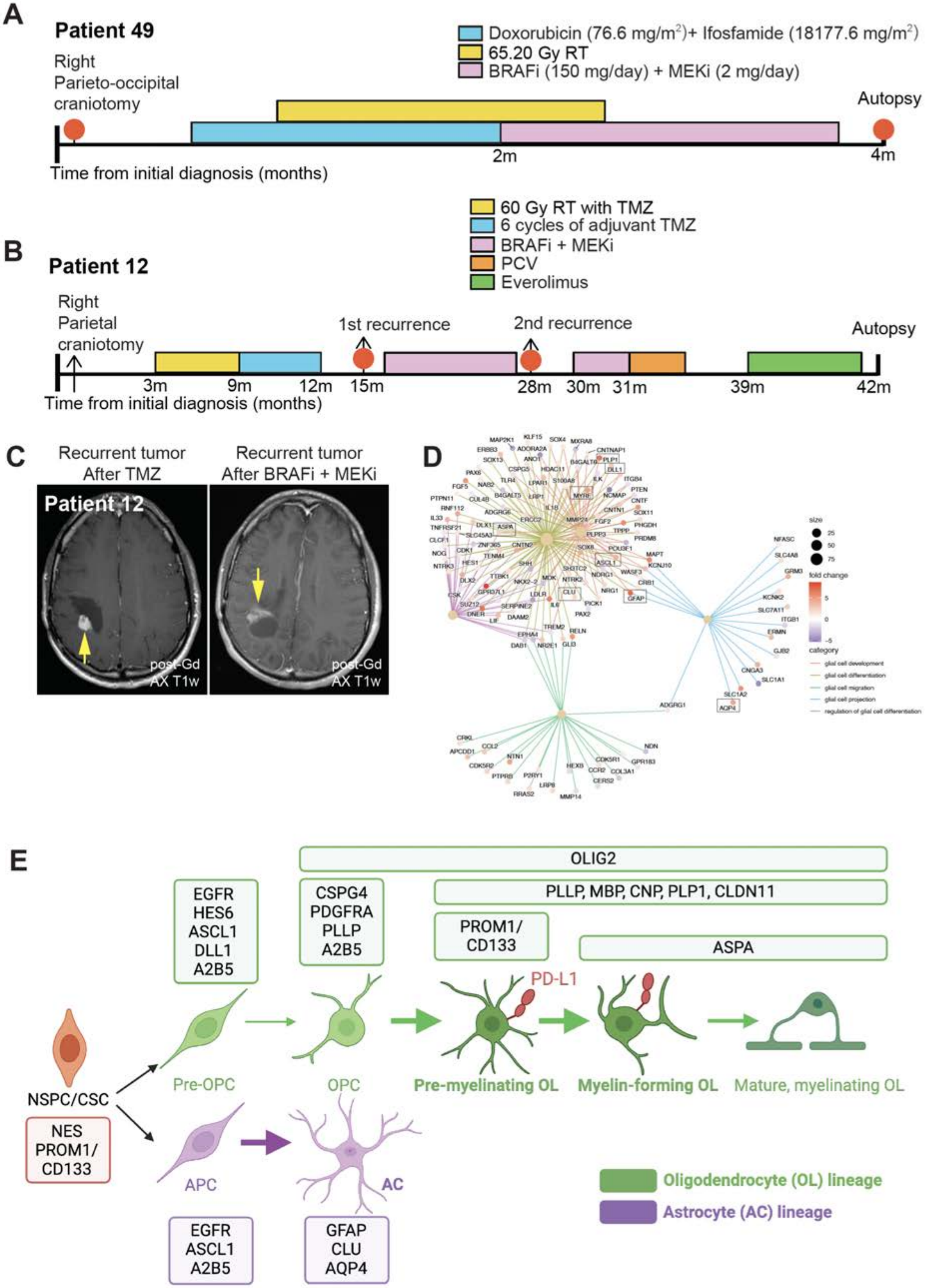

A
Patient 49
Doxorubicin (76.6 mg/m2)+ Ifosfamide (18177.6 mg/m2)
65.20 Gy RT
BRAFi (150 mg/day) + MEKi (2 mg/day)
Right Parieto-occipital craniotomy
Autopsy
2m
4m
Time from initial diagnosis (months)
B
Patient 12
60 Gy RT with TMZ
6 cycles of adjuvant TMZ
BRAFi + MEKi
PCV
Everolimus
Right Parietal craniotomy
1st recurrence
2nd recurrence
Autopsy
3m
9m
12m
15m
28m
30m
31m
39m
42m
Time from initial diagnosis (months)
C
Recurrent tumor After TMZ
Recurrent tumor After BRAFi + MEKi
Patient 12
post-Gd AX T1w
post-Gd AX T1w
D
E
OLIG2
PLLP, MBP, CNP, PLP1, CLDN11
PROM1/ CD133
ASPA
EGFR HES6 ASCL1 DLL1 A2B5
CSPG4 PDGFRA PLLP A2B5
PD-L1
NSPC/CSC
NES PROM1/ CD133
Pre-OPC
OPC
Pre-myelinating OL
Myelin-forming OL
Mature, myelinating OL
APC
AC
EGFR ASCL1 A2B5
GFAP CLU AQP4
Oligodendrocyte (OL) lineage
Astrocyte (AC) lineage

**Figure S1. BRAF$^{V600E}$-mutant HGG Undergo Cell State Transitions Along Glial Differentiation Trajectories Following Extensive Treatment in Patients.**

**(A-B)** A timeline of treatment histories for patient 49 (A) and patient 12 (B) with BRAF$^{V600E}$ mutant high-grade glioma (HGG) at the time of diagnosis. Red circles correspond to the time of recurrent tumor tissue collection for subsequent analyses (corresponding to Figure. 1A). **(C)** Magnetic resonance imaging (MRI) contrast-enhanced images axial view revealed tumor lesion (yellow arrow) at the 1$^{st}$ recurrence and 2$^{nd}$ recurrence post-treatment with BRAFi+MEKi in patient 12. **(D)** The cnetplot showing specific genes enriched in gliogenesis-associated pathways from the gene ontology analysis at 2$^{nd}$ tumor recurrence after BRAFi+MEKi treatment in patient 12. **(E)** Schematic of glial differentiation trajectories with cell state-specific markers indicated in the boxes. NSPC: neural stem/progenitor cell; CSC: cancer stem cell; Pre-OPC: pre-oligodendrocyte progenitor cell; APC: astrocyte progenitor cell; OL: oligodendrocyte; AC: astrocyte. Related to Figure 1.

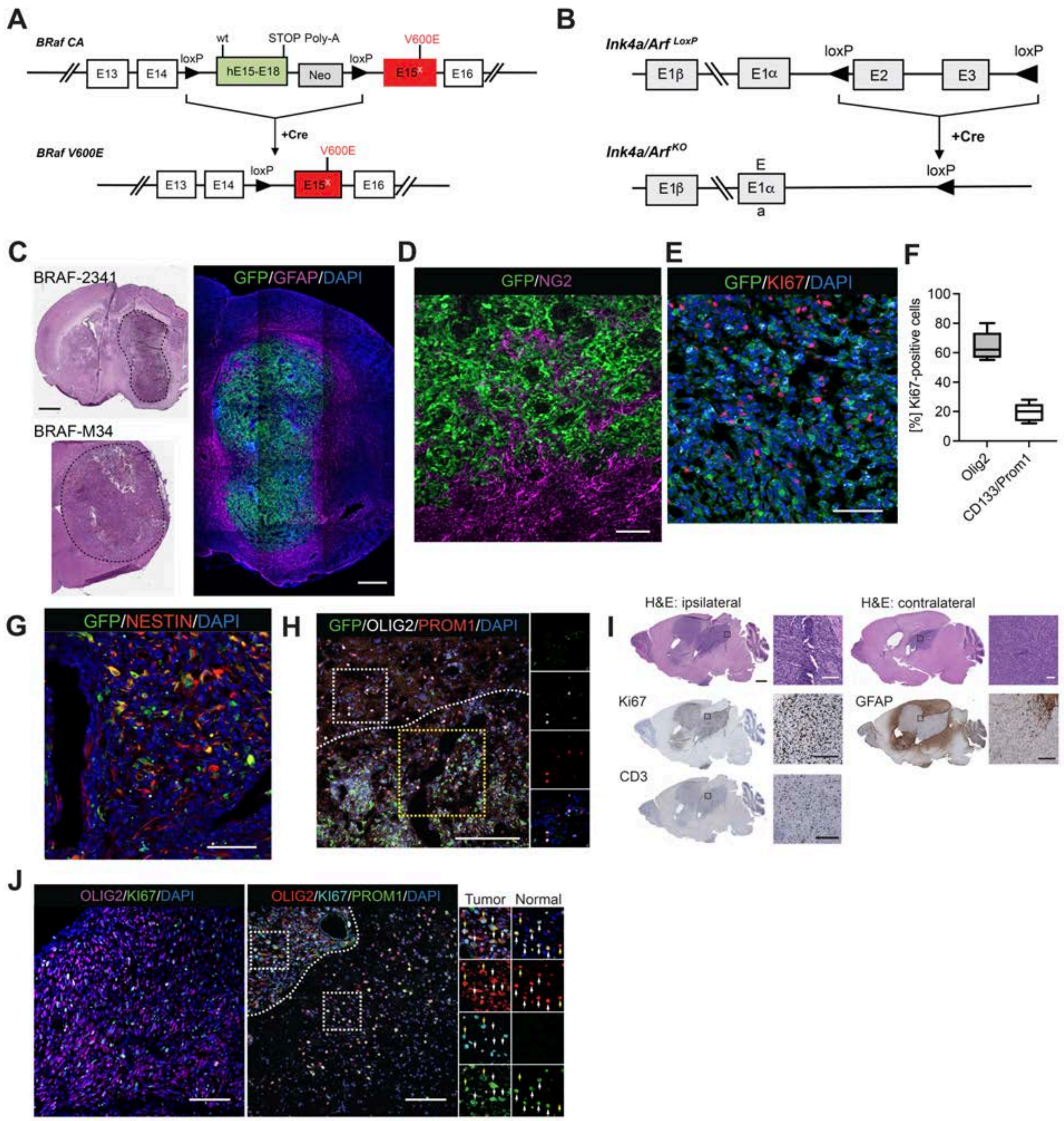


**Figure S2. Novel Orthotopic BRAF$^{V600E}$-mutant Mouse Models Show Intra-tumoral Cellular Heterogeneity.**

**(A)** Schematic diagram of the Braf cre-activatable (CA) knock-in allele in the absence (top) and presence of Cre expression (bottom). **(B)** Schematic diagram of the Ink4a/Arf knock-out allele in the absence (top) and presence of Cre expression (bottom). **(C)** Left panel: Representative images of H&E-stained coronal mouse brain sections of BRAF-2341 and BRAF-M34 models revealed the formation of orthotopic tumors upon BRAF V600E expression and deletion of *Cdkn2a*. Right panel: an IF image of the entire BRAF-2341 mouse brain section stained for GFP, GFAP, and DAPI for DNA. Scale bars: 1 mm (left), 520µm (right). **(D)** Representative IF image of BRAF-2341 HGG stained for GFP and NG2. Scale bars: 70µm. **(E)** Representative IF image of BRAF-2341 HGG stained for GFP, Ki67, and DAPI for DNA. Scale bar: 100µm. **(F)** Quantification revealed that the percentage of CD133/PROM1-positive cells amongst Ki67-

positive cells (19.4%) was significantly lower than that of Olig2-positive cells (64.4%). ****$P<0.0001$, unpaired t-test. Five individual tumors were counted. **(G)** Representative IF image of BRAF-2341 HGG stained for GFP, NESTIN, and DAPI for DNA. Scale bar: 100μm. **(H)** Representative IF images of BRAF-2341 HGG (below the dashed line) and normal cells (above the dashed line) stained for GFP, OLIG2, PROM1, and DAPI for DNA. Single-channel images of the white-boxed area are shown next to the merged image. The merged image of the yellow boxed area is shown in Figure 2D, middle panel. Scale bar: 160μm. **(I)** Representative images of H&E and immunohistochemically stained sagittal sections of RCAS-BRAF glioma showing large glioma mass in the ipsilateral and contralateral hemispheres, indicative of infiltrative growth. Scale bars: 1 mm (low magnification), 100μm (high magnification). **(J)** Representative IF images of RCAS-BRAF HGG stained for OLIG2, Ki67, PROM1, and DAPI for DNA. Single-channel images of the white boxed areas above (tumor cells) and below (normal cells) the dashed line are shown next to the merged image. Tumor cells are either Olig2/Ki67/Prom1 triple-positive (yellow arrows), Olig2/Prom1 double-positive (white arrows), or Olig2 positive only (white asterisks). Normal oligodendroglial cells negative for Ki67 are either Olig2/Prom1 double-positive (white arrows) or Olig2 positive only (yellow arrows). Scale bars=140μm (left), 130μm (right). Related to Figure 2.

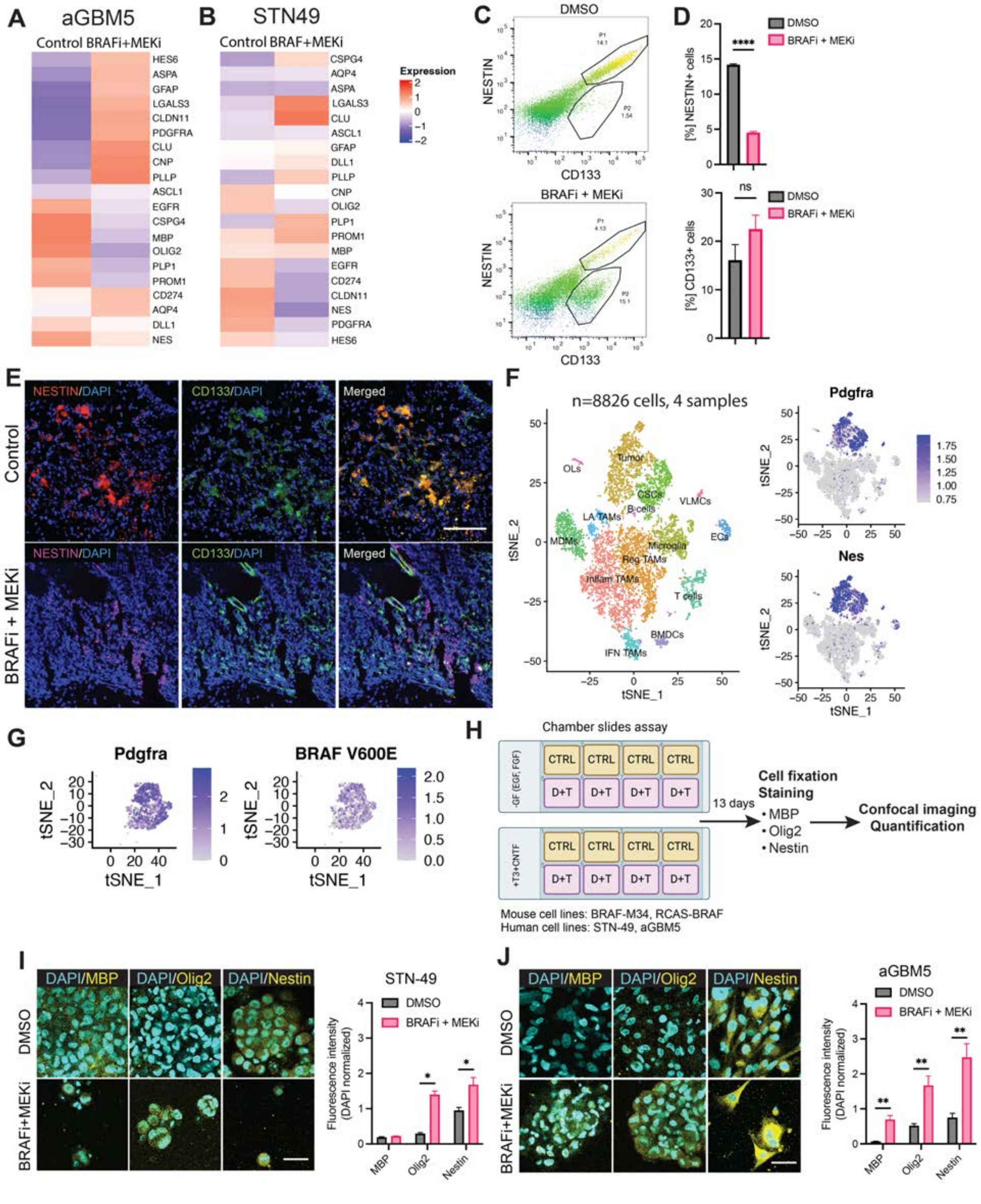


**Figure S3. BRAFi+MEKi Treatment Heightens Cell State Transitions Along Glial Differentiation Trajectories in BRAF$^{V600E}$-mutant HGG.**

**(A-B)** Average expression levels of genes of interest associated with stem/progenitor cells and glial cell differentiation between two replicates using patient-derived SU-aGBM5 (A) and STN-49 (B) cell lines treated with BRAFi+MEKi for 48 hours. **(C)** Representative flow cytometry dot plots for BRAF-2341 HGG cells stained for PROM1/CD133 and NESTIN. Top: Control-treated cells (DMSO). Bottom: BRAFi+MEKi-treated cells. P1=PROM1/CD133 and NESTIN double positive. P2=PROM1/CD133-positive, NESTIN-negative. **(D)** Quantification of flow cytometry analysis from (C). ****P<0.0001, unpaired t-test. Ns, not significant. **(E)** Representative IF images of BRAF-2341 HGG stained for NESTIN, CD133, and DAPI for DNA. Scale bar: 100µm. **(F)** tSNE plot revealing cell clusters of the integrated dataset of all samples obtained from the RCAS-BRAF model using unsupervised clustering in Seurat. BMDCs: Bone marrow-derived dendritic cells; CSCs: Cancer stem cells; ECs: Endothelial cells; IFN TAMs: IFNg-primed tumor-associated macrophages (TAMs); Inflam TAMs: inflammatory TAMs; LA TAMs: lipid-associated TAMs; MDMs: Monocyte-derived macrophages; OLs: oligodendrocytes; Reg TAMs: regulatory TAMs; VLMCs: Vascular leptomeningeal cells. Feature plots on the right show tumor/CSC-specific markers (*Pdgfra, Nes*) from the RCAS-BRAF dataset. The expression of genes is indicated by a color scale ranging from gray (no expression) to dark blue (high expression). **(G)** Feature plots showing BRAF V600E expression in the subclustered Pdgfra-positive tumor/CSCs from the RCAS-BRAF dataset. The expression of genes is indicated by a color scale ranging from gray (no expression) to dark blue (high expression). **(H)** Schematic of the experimental design for assessing the capacity of BRAFi+MEKi to promote tumor cell differentiation *in vitro*. **(I-J)** Representative ICC images of patient-derived STN-10049 (STN-49) (I) and SU-aGBM5 (aGBM5) (J) cell lines in spontaneous differentiation medium (except for aGBM5 which was incubated with differentiation medium) when treated with vehicle and concomitant BRAFi+MEKi stained for MBP, OLIG2, NESTIN and DAPI for DNA. *P<0.05, **P<0.01 by unpaired multiple t-tests and Holm-Šídák method. n=2 independent experiments, data represented as mean±SEM. Related to Figure 3.

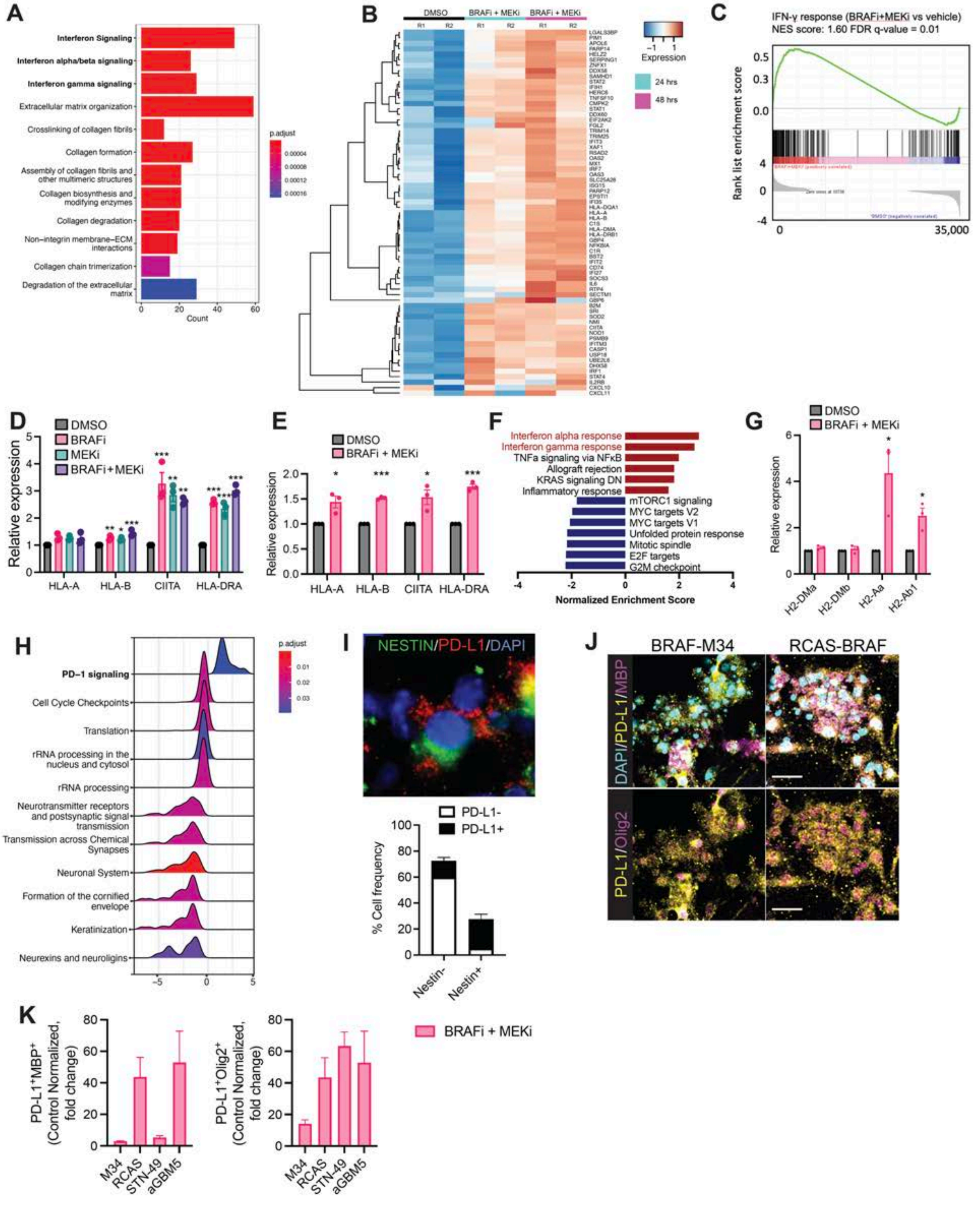

**Figure S4. BRAFi+MEKi Treatment Induces Interferon Response Pathway Gene Expression Signature and Glial Differentiation Cell State Transitions, with a PD-L1 expressing immunomodulatory subpopulation.**

**(A)** Differential expression analysis and gene set enrichment analysis (GSEA) of gene expression data from the patient-derived aGBM5 cell line revealed significant upregulation of genes associated with interferon signaling after 48 hours of BRAFi+MEKi treatment. **(B)** Heatmap demonstrating significant increases in expression of IFN-γ receptor-associated transcripts after BRAFi+MEKi treatment compared with vehicle treatment at 24 and 48 hours in the aGBM5 cell line. R: technical replicate. **(C)** GSEA enrichment plot of gene expression data from aGBM5 cell line comparing BRAFi+MEKi treatment with vehicle treatment. NES, normalized enrichment score. FDR: false discovery rate. **(D-E)** Validation of specific MHC gene upregulation by qRT-PCR in aGBM5 (D) and STN-49 (E) cell lines. Data shown from 48 hours of BRAFi+MEKi treatment compared with DMSO control treatment were assessed as mean ± SEM of at least 3 independent experiments. $^{*}P < 0.05$; $^{**}P < 0.01$; $^{***}P < 0.001$ by unpaired t-test. **(F)** Validation of interferon response as the top upregulated gene expression pathway post-BRAFi+MEKi treatment in acutely isolated BRAF-2341 cells. Two-sided bar chart showing the normalized GSEA enrichment score. **(G)** Relative expression of MHC genes post-BRAFi+MEKi treatment in BRAF-2341 HGG cells at 48 hours of treatment assessed by qRT-PCR. H2-Aa (synonym-HLA IA-alpha) and H2-Ab1 (synonym-HLA IA-beta) were significantly upregulated. Data are shown as mean ± SEM of at least 3 independent experiments. $^{*}P < 0.05$ by unpaired t-test. **(H)** Ridge plot of significantly upregulated Reactome pathway “PD-1 signaling” in aGBM5 cell line after BRAFi+MEKi treatment for 48 hrs. **(I)** Representative IF image of BRAF-2341 HGG stained for PD-L1, NESTIN and DAPI for DNA. n=3 mice/group. Quantification of (I). **(J)** Representative ICC images of murine cells treated with BRAFi+MEKi for 13 days, and subsequently stained for PD-L1, MBP, OLIG2 and DAPI for DNA. Scale bars: 40μm. **(K)** Relative cell number (fold changes) of MBP or OLIG2 and PD-L1 double-positive cells in murine and human cell lines treated with BRAFi+MEKi for 13 days. Related to Figure 4.

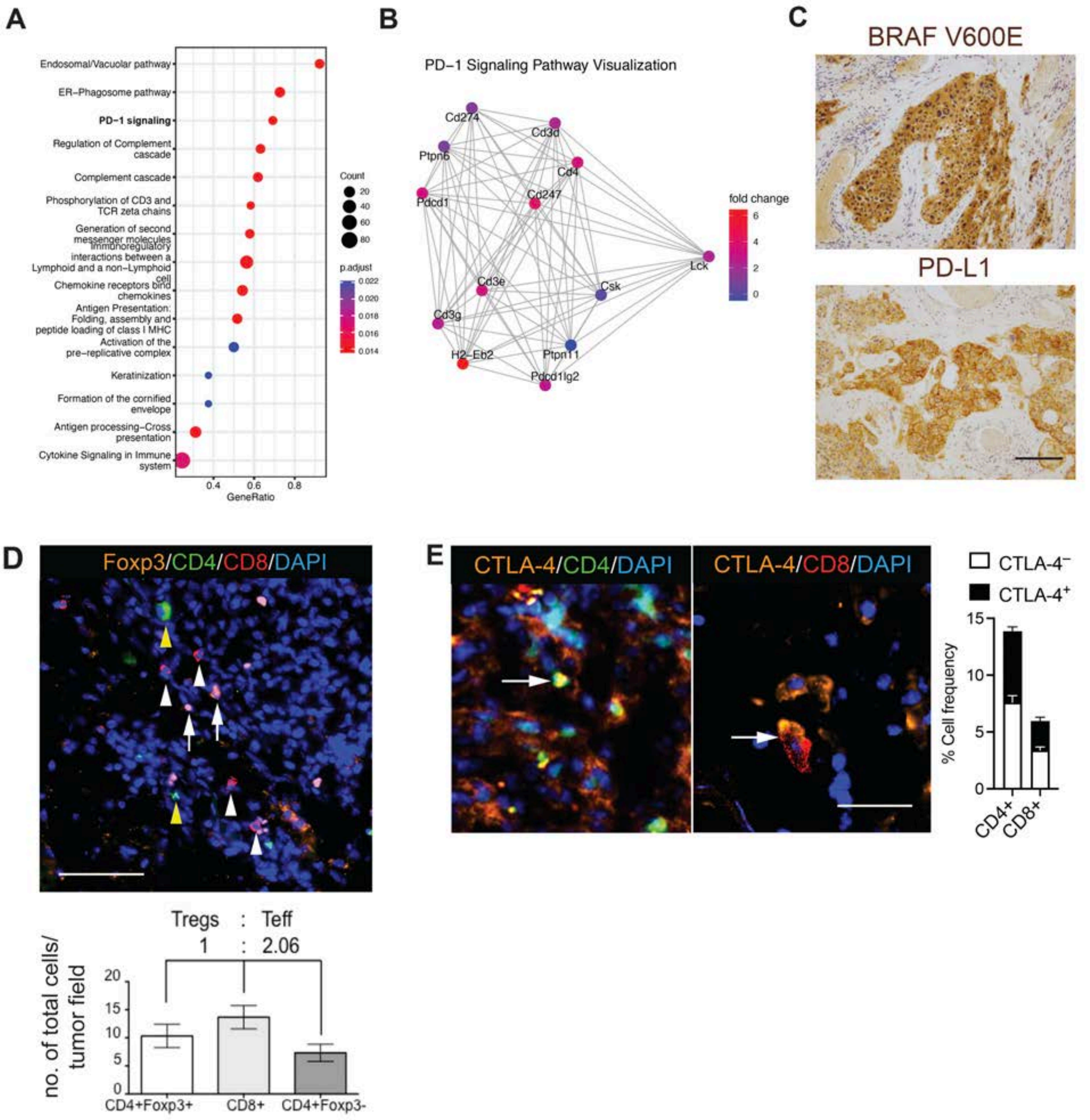


**Figure S5. BRAFi+MEKi Treatment Induces PD-L1 Expression and T Cell Infiltration in Murine BRAF$^{V600E}$-mutant HGG.**

**(A)** PD-1 signaling (bold) is the third top significantly upregulated Reactome pathway post-BRAFi+MEKi for 72hrs as shown in a dotplot in BRAF-2341 cells. **(B)** Visualization of PD-1 signaling network in BRAF-2341 cells post-BRAFi+MEKi for 72 hours. **(C)** Representative IHC images for BRAF V600E and PD-L1 in BRAF$^{V600E}$-mutant HGG. Scale bar: 25µm. **(D)** Representative IF images of BRAF-2341 HGG stained for Foxp3, CD4, CD8 and DAPI for DNA. Yellow arrowheads: CD4+ T cells, white arrowheads: CD8+ T cells, white arrows: CD4+Foxp3+ regulatory T cells. The quantification of the cell populations is shown on the bottom. Scale bar: 20µm. **(E)** Representative IF images of BRAF-2341 HGG stained for CTLA-4, CD4, CD8, and DAPI for DNA. Arrows indicate CD4+ and CD8+ cells co-expressing CTLA-4. Quantification of cell populations. n=3 mice/group. Scale bar, 5µm. Related to Figures 4, 5 and 6.

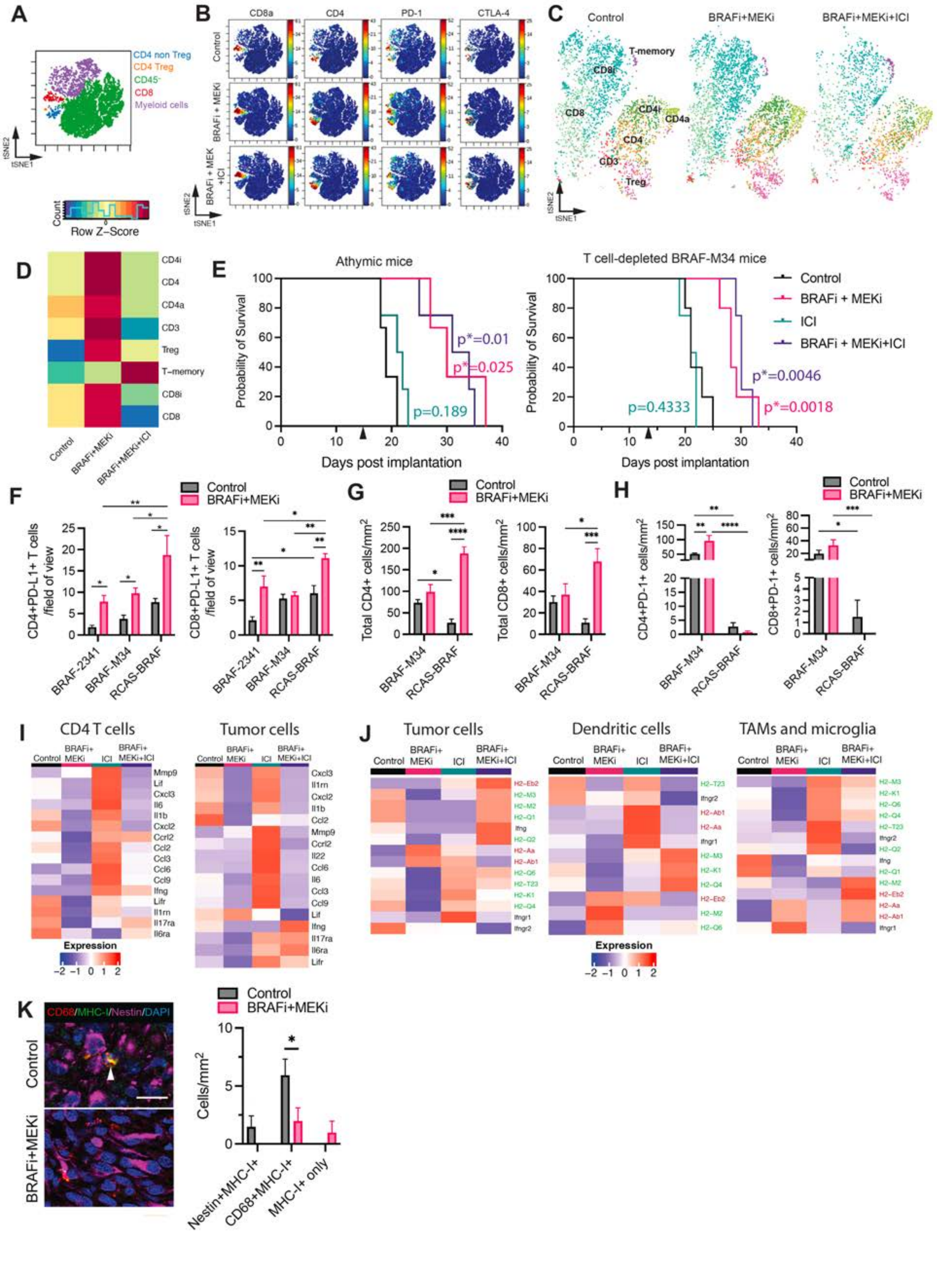
A
CD4 non Treg
CD4 Treg
CD45-
CD8
Myeloid cells
tSNE2
tSNE1
Row Z-Score
B
CD8a
CD4
PD-1
CTLA-4
Control
BRAFi + MEKi
BRAFi + MEK +ICI
C
Control
BRAFi+MEKi
BRAFi+MEKi+ICI
T-memory
CD8i
CD8
CD4i
CD4a
CD4
CD3
Treg
D
CD4i
CD4
CD4a
CD3
Treg
T-memory
CD8i
CD8
Control
BRAFi+MEKi
BRAFi+MEKi+ICI
E
Athymic mice
T cell-depleted BRAF-M34 mice
Probability of Survival
Days post implantation
p*=0.01
p*=0.025
p=0.189
p=0.4333
p*=0.0046
p*=0.0018
Control
BRAFi + MEKi
ICI
BRAFi + MEKi+ICI
F
Control
BRAFi+MEKi
CD4+PD-L1+ T cells /field of view
CD8+PD-L1+ T cells /field of view
BRAF-2341
BRAF-M34
RCAS-BRAF
G
Total CD4+ cells/mm2
Total CD8+ cells/mm2
H
CD4+PD-1+ cells/mm2
CD8+PD-1+ cells/mm2
I
CD4 T cells
Tumor cells
Control
BRAFi+ MEKi
ICI
BRAFi+ MEKi+ICI
Expression
J
Tumor cells
Dendritic cells
TAMs and microglia
K
CD68/MHC-I/Nestin/DAPI
Control
BRAFi+MEKi
Cells/mm2
Nestin+MHC-I+
CD68+MHC-I+
MHC-I+ only

**Figure S6. BRAFi+MEKi Enhances T Cell Infiltration and Synergizes with ICI to Improve T Cell-Dependent Survival.**

**(A)** Representative t-distributed stochastic neighbor embedding (tSNE) map of cell populations from CyTOF analysis. **(B)** Representative CyTOF tSNE plots depicting the relative expression levels of CD8a, CD4, PD-1, and CTLA-4 of BRAF-2341 HGG from the control and treatment groups. **(C)** Representative CyTOF tSNE plots depicting the relative expression levels of CD8i, CD8, CD4a, CD4i, CD4, CD3, Treg, and T-memory cells of syngeneic, orthotopic BRAF-2341 mice from the control and treatment groups. **(D)** Heatmap showing the relative expression levels of CD8i, CD8, CD4a, CD4i, CD4, CD3, Treg, and T-memory cells of syngeneic, orthotopic BRAF-2341 mice from the control and treatment groups. **(E)** Kaplan-Meier survival curve in immunocompromised (athymic) mice with orthotopic BRAF-2341 tumor cells (left panel) or T cell-depleted C57BL/6 mice with orthotopic BRAF-M34 tumor cells (right panel) treated with BRAFi+MEKi and/or ICI. T-cell depletion was induced by injecting mice with α-CD4 and α-CD8 antibodies every four days throughout the experiment. Black arrows indicate the treatment initiation at day 14 post-injection. Data is representative of two independent experiments. 5-6 mice/group (athymic) and 4-5 mice/group (T cell-depleted mice). P values are indicated in the graph and Tables 1 and 2 when compared to the control group. **(F)** IF quantification of CD4+ and CD8+ T cells immunoreactive for PD-L1 in BRAF-2341, BRAF-M34, and RCAS-BRAF HGGs before versus after BRAFi+MEKi treatment. $^*P<0.05$, $^{**}P<0.01$ by two-way ANOVA with Sidak's multiple comparisons test. n=4-5 mice/group, data represented as mean±SEM. **(G)** IF quantification of total CD4+ and CD8+ T cells in BRAF-M34, and RCAS-BRAF HGGs before versus after BRAFi+MEKi treatment. $^*P<0.05$, $^{***}P<0.001$, $^{****}P<0.0001$ by two-way ANOVA with uncorrected Fisher's LSD. n=4 mice/group, data represented as mean±SEM. **(H)** IF quantification of CD4+ and CD8+ T cells immunoreactive for PD-1 in BRAF-M34 and RCAS-BRAF HGGs before versus after BRAFi+MEKi treatment. $^*P<0.05$, $^{**}P<0.01$, $^{***}P<0.001$, $^{****}P<0.0001$ by two-way ANOVA with uncorrected Fisher's LSD. n=4 mice/group, data represented as mean±SEM. **(I)** Heatmaps of selected pro-inflammatory cytokines in CD4 T cells and tumor cells in the RCAS-BRAF model with different treatment groups. **(J)** Heatmaps of selected MHC class I (green), MHC class II (red), and IFNg-related genes (black) associated with antigen presentation in tumor cells, dendritic cells, and tumor-associated macrophages (TAMs) and microglia in orthotopic RCAS-BRAF HGG with different treatment groups. **(K)** Representative IF images of RCAS-BRAF tissue stained for NESTIN, CD68, MHC-I, and DAPI. The arrow indicates CD68 and MHC-I double-positive cells. Quantification of cell populations after BRAFi+MEKi treatment compared with vehicle control treatment. $^*P < 0.05$ as determined by two-way ANOVA with Sidak's multiple comparisons test. n=4 mice per group, data represented as mean±SEM. Scale bar: 15μm. Related to Figure 6.

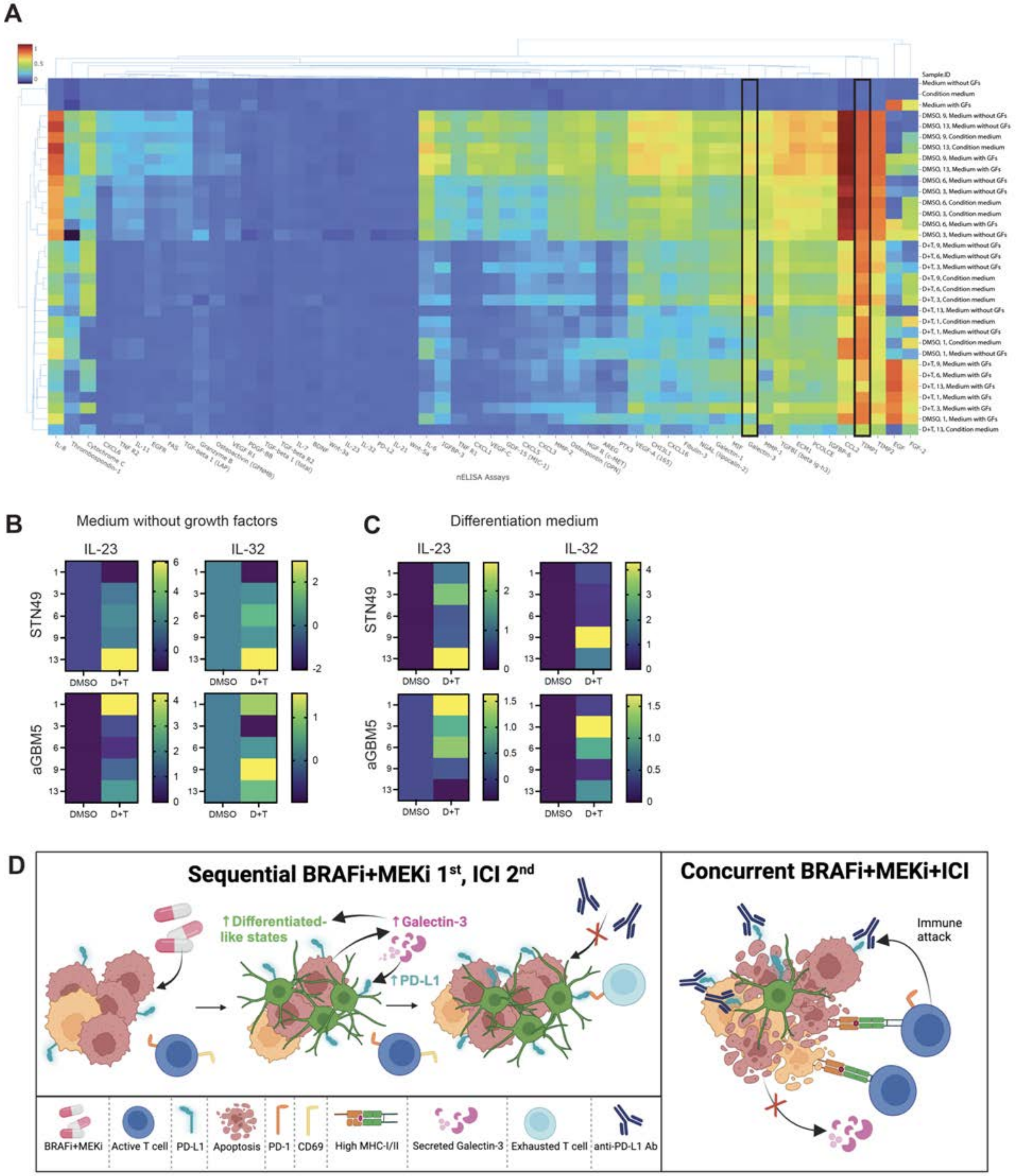


**Figure S7. BRAFi+MEKi-Induced Secretion of Galectin-3, IL-23 and IL-32 From Tumor Cells that Could Play Roles in Immunosuppression.**

**(A)** Clustergram representation of normalized nELISA signals from selected secreted markers. **(B-C)** Relative expression levels of IL-23 and IL-32 in medium deprived of growth factors (B) or in differentiation medium (C). The color bar represents normalized expression values, which are presented as z-scores. **(D)** Hypothetical model of mechanisms between sequential vs. concurrent quadruple treatment. Related to Figure 7.

**Table S1. Clinical characteristics of two patients with BRAF$^{V600E}$-mutant high-grade glioma at the time of diagnosis.**

| Patient ID | Histopath Diagnosis | Sex | Age at Diagnosis | Exposure Time (mo) | Best Response | Time to recurrence (mo) |
|---|---|---|---|---|---|---|
| **Patient 12** | HGG/ GBM | Male | 22 | 13 | Stable Disease | 15 |
| **Patient 49** | HGG/ GBM | Male | 13 | 3 | Progressive Disease | 4 |

**Table S2. A summary table of histological, immunophenotypic, and pharmacological analyses of tumor cells in novel orthotopic, syngeneic mouse models for BRAF$^{V600E}$-mutant high-grade gliomas.**

| Models | CRE-BRAF-2341 (BRAF-2341) | CRE-BRAF-M34 (BRAF-M34) | RCAS-BRAF |
|---|---|---|---|
| **Genetic background** | FVB/N | C57BL/6 | C57BL/6 |
| **Number of cells injected** | 250k in 2.5µl | 250k in 2.5µl | 50k in 1µl |
| **Genotype** | BRAF V600E het CDKN2A del hmz | BRAF V600E het CDKN2A del hmz | RCAS-BRAF V600E TP53 del (shRNA) |
| **Region** | Striatum | Striatum | Striatum |
| **MRI** | diffuse, extra- and intra-parenchymal growths | strong rim enhancement and necrotic centers | infiltrative |
| **Necrosis** | High | High | Low |
| **Microvascular proliferation** | High | High | Moderate |
| **Nuclear pleomorphism** | High | High | High |
| **Grade** | HGG | HGG (PXA) | HGG |
| **Marker expression** | Nestin +<br>Olig2 ++<br>GFAP ++<br>PROM1 ++<br>CSPG4 + | Nestin +<br>Olig2 +<br>PDGFRa ++<br>GFAP +<br>NeuN +++ | Nestin +++<br>Olig2 ++<br>PDGFRa +++<br>GFAP +<br>NeuN ++ |
| **BRAFi+MEKi response** | Sensitive | Sensitive | Resistant |

**Table S3. Oligonucleotide sequences for RT-qPCR**

| | |
|---|---|
| GAPDH_F | TGGGGAAGGTGAAGGTCGG |
| GAPDH_R | CTGGAAGATGGTGATGGGA |
| HLA-A_F | AAAAGGAGGGAGTTACACTCAGG |
| HLA-A_R | GCTGTGAGGGACACATCAGAG |
| HLA-B_F | CTACCCTGCGGAGATCA |
| HLA-B_R | ACAGCCAGGCCAGCAACA |
| HLA-DRA_F | GCCAACCTGGAAATCATGACA |
| HLA-DRA_R | AGGGCTGTTCGTGAGCACA |
| CIITA_F | GGCTGGAATTTGGCAGCAC |
| CIITA_R | GCCCAACACAAGGATGTCTCT |
| Gapdh_F | CAGTATGACTCCACTCACGGCA |
| Gapdh_R | CCTTCTCCATGGTGGTGAAGAC |
| IA alpha_F | CTTCCCACCTGTGATCAACAT |
| IA alpha_R | AATCTCAGGTTCCCAGTGTTT |
| IA beta_F | GACGCAGCGCATACGGCTC |
| IA beta_R | CCGCCGCAGGGAGGTGCT |

**Table S4. List of patient tumors for PD-L1 analysis.**

| Patient ID | PD-L1 Frequency Score (%) | PD-L1 Intensity Score (%) | PD-L1 Score | Diagnosis |
|---|---|---|---|---|
| **E-338/2017** | 95 | moderate | 8 | Ganglioglioma |
| **E-1794/2017-2** | 20 | weak | 2 | Ganglioglioma |
| **E-1869/2014** | 70 | moderate | 8 | PXA |
| **E-831/2017** | 90 | strong | 12 | Malignant glioma |
| **E-2377/2015** | 65 | moderate | 8 | Malignant astrocytoma |
| **E-2012/2014** | 50 | weak | 4 | PXA |
