## Supplementary material for "BRAF/MEK Inhibition Induces Cell State Transitions Boosting Immune Checkpoint Sensitivity in BRAF^V600E^-mutant Glioma": Methods

### Key Resources Table

See Key Resources Table

### Resources Availability

#### Lead Contact

#### Materials availability

Murine BRAF-M34, RCAS-BRAF cells, and patient-derived cell lines will be made available upon request and require a materials transfer agreement according to Stanford guidelines.

#### Data and code availability

Data supporting the findings of this study are available in the article and its supplementary files. Bulk RNA sequencing data have been deposited in NCBI's Gene Expression Omnibus (GEO) under accession number GSE287972 [public release upon manuscript acceptance]. Single-cell RNA sequencing data have been deposited in GEO and are accessible through accession number GSE288791 [public release upon manuscript acceptance]. The codes for analyzing bulk and single-cell RNA-seq data; as well as NOMIC nELISA and CYTOF data, are available upon request. All other datasets not included in this publication are available from the corresponding author upon reasonable request. Links of all used tools are listed in the Key Resources Table.

### Experimental Model and Subject Details

#### Animal Studies

For mouse studies male and female mice are used. All animal procedures were performed in accordance with institutional and federal guidelines (Stanford University APLAC protocol # 33633). Mice were housed in groups of three to five and maintained on a 12-hour light/dark schedule. Food and water were available *ad libitum*. Compound transgenic mice were generated by crossing Cre-dependent BRAF CA with Ink4a/Arf flox/flox mice. *BRAF* $^{CA/WT}$*; Ink4a/Arf* $^{flox/flox}$ mice were of a pure FVB/N or C57BL/6 genetic background. Littermates carrying appropriate genotype(s) were randomly assigned to experimental groups. Mice were intracranially injected postnatally or at 6-8 weeks of age. Mice were treated and analyzed at specific postnatal days or in adulthood, as indicated in the text and figure captions.

#### Human Studies

Tumor specimens and clinical information were collected according to protocols approved by the Research Ethics Board. Patients provided informed consent, and ethical approval was obtained from the Institutional Review Board at Stanford University. Formalin-fixed, paraffin-embedded (FFPE) brain tumor tissue was obtained from patients with $BRAF^{V600E}$-mutant gliomas. Clinically relevant dual MAPK pathway inhibitors were administered to a 13-year-old boy with $BRAF^{V600E}$-mutant high-grade GBM, who eventually succumbed to the disease. The patient was initially treated with one cycle of doxorubicin/ifosfamide treatment with 65.20 Gy in 35 fractions of radiotherapy. The patient subsequently underwent dabrafenib and trametinib treatment until his death. Separate primary patient samples were also obtained from Children's Hospital Colorado (Aurora, CO) in accordance with local and federal human research protection guidelines and Institutional Review Board regulations. Clinical and pathological features of patient 12 with $BRAF^{V600E}$-mutant glioma were described in Schreck et al., 2021. Specifically, this young adult man diagnosed with high-grade GBM was initially treated with temozolomide with 60 Gy in 30 fractions of radiotherapy. The patient subsequently received 6 cycles of adjuvant temozolomide for 6 months. However, given his first local recurrence 3 months later, this patient then underwent dabrafenib and trametinib treatment for 13 months. All tumor specimens were given a sequential unique identifier to ensure complete deidentification.

PD-L1 Expression:

For PD-L1 expression comparisons in adult GBM, clinical pathology data were obtained from 3126 glioblastoma tissue specimens profiled at Caris® Life Sciences. All samples were from adult patients (≥18 years of age). Each tumor was reviewed by board-certified pathologists and classified as a glioblastoma based on World Health Organization (WHO) classification guidelines. Given changes in the WHO classification of GBM, samples with mutations in *IDH1* or *IDH2* were excluded from analyses to conform with 2021 revisions to the WHO classification of brain tumors. Tumor specimens were collected via manual microdissection and harvesting targeted tumor tissue. Next-generation sequencing was performed on genomic DNA isolated from FFPE samples using either a custom-designed SureSelect XT assay (592 whole-gene targets, Agilent Technologies, Santa Clara, CA) or whole-exome sequencing. All variants were detected with >99% confidence based on allele frequency and amplicon coverage, with an average sequencing depth across all amplicons >500x. DNA sequencing results are validated per Clinical Laboratory Improvement Amendments (CLIA) and Internal Organization for Standardization (ISO) requirements. *MGMT*p pyrosequencing and IHC for PD-L1 (SP142 clone) were performed on all samples. ArcherPlex Fusion profiling data were available for 69% of samples, and bulk whole-transcriptome sequencing data were available for 33% of samples.

For comparisons in Fig 5A, PD-L1 expression was tested by IHC using SP142 antibody (Spring Biosciences). The staining intensity on the tumor cells was assessed on a semiquantitative scale: 0 for no staining, 1+ for weak staining, 2+ for moderate staining, and 3+ for strong staining. Tumors exhibiting ≥5% of tumor cells stained as 2+ or 3+ were regarded as being PD-L1 positive, as previously described (Arai, Hiroyuki et al. "Molecular Characterization of Appendiceal Goblet Cell Carcinoid." *Molecular cancer therapeutics* vol. 19,12 (2020): 2634-2640. doi:10.1158/1535-7163.MCT-20-0318).

For comparisons in Fig 5B, the number of *CD274* (ie, PD-L1) transcripts per million in the BRAF mutation/fusion group (n=17) were compared to the number in the BRAF-WT group (n=1022) using a Wilcoxon Rank-Sum test. This non-parametric test does not assume normality of the CD274 transcript counts, and has been shown to demonstrate asymptotic normality and be robust in the setting of small and unbalanced sample sizes. Specifically, the test tends to be conservative in the setting of unbalanced samples but retains greater statistical power than other common alternatives (e.g., median tests). See reference: Mann, H. B., & Whitney, D. R. (1947). On a test of whether one of two random variables is stochastically larger than the other. *Annals of Mathematical Statistics, 18,* 50–60. https://doi.org/10.1214/aoms/1177730491.

### Cell Lines and Primary Cultures

All cell lines and primary cultures were maintained in humidified cell culture incubators at 37$^o$C under 5% $CO_2$. SU-aGBM5 is a BRAF$^{V600E}$ mutant GBM cell line from a young adult obtained from Dr. Michelle Monje (Stanford University, CA, USA). Murine BRAF$^{V600E}$ mutant glioma cell lines (BRAF-2341, BRAF-M34) were derived from murine glioma, induced by injection of adenovirus-Cre into the brain of *BRAF* $^{CA/WT}$*;Ink4a/Arf* $^{flox/flox}$ mice. Cells were maintained in 50% Neurobasal-A medium without L-Glutamine (10888022; ThermoFisher Scientific, Waltham, MA) and 50% D-MEM/F12 medium (11330-032; ThermoFisher Scientific), supplemented with 0.1mM MEM Non-Essential Amino Acids Solution (11140-050; ThermoFisher Scientific), 1 mM MEM Sodium Pyruvate Solution (11360-070; ThermoFisher Scientific), 1X GlutaMAX-I Supplement (35050-061; ThermoFisher Scientific) and 10 mM HEPES Buffer (15630-080; ThermoFisher Scientific). Gibco B27 Supplement without vitamin A (12587010; ThermoFisher Scientific) and N2 supplement (17502048; ThermoFisher Scientific) were also added to the medium. Cells were supplemented twice a week with 40 ng/ml FGF (100-146; Fujifilm Irvine Scientific, Santa Ana, CA) and EGF (100-26; Fujifilm Irvine Scientific), and 10 ng/ml PDGF-AA (100-16; Fujifilm Irvine Scientific), PDGF-BB (100-18; Fujifilm Irvine Scientific) and STEMCELL Technologies heparin (7980; ThermoFisher Scientific). BRAF-M34, RCAS-BRAF, and aGBM5 cell lines were cultured on laminin (L2020; Sigma-Aldrich, St. Louis, MO), while STN49 grew on pol-l-ornithine (P3655; Sigma-Aldrich), following manufacturer's instructions.

### Intracranial Injections of Adenovirus

*BRAF*$^{CA/WT}$*; Ink4a/Arf* $^{flox/flox}$ mice were injected with 2μl of adenovirus expressing cre recombinase under the control of the ubiquitous CMV promoter (Ad-Cre; Titer = 1x10^10 PFU/ml; Vector Biolabs, Malvern, PA) to induce BRAF V600E expression and deletion of tumor suppressor genes p16 $^{Ink}$ and p19 $^{Arf}$ (Ink4a/Arf: murine Cdkn2a homologous locus). Control mice were injected with adenovirus expressing GFP (Ad-GFP; Vector Biolabs). The stereotaxic coordinates selected to target the cortical layer, relative to bregma, are anterior-posterior–1.00 mm, medial-lateral–1.80 mm, and dorsal-ventral–1.5 mm. Animals were observed and sacrificed when exhibiting clinical symptoms consistent with intracranial tumor formation. Brains of symptomatic mice were resected, and sectioned and tumor-bearing brains were subjected to hematoxylin and eosin (H&E) and immunofluorescence staining.

**Orthotopic Glioma Generation**

6-8-week-old FVB/N and nu/nu mice were used for orthotopic implantation of BRAF-2341 glioma cells, while age-matched C57BL/6 mice were used for orthotopic implantation of BRAF-M34 glioma cells. C57BL/6 mice at postnatal day 7 were used for orthotopic implantation of RCAS-BRAF murine glioma cells. Adult and postnatal mice were anesthetized with isoflurane inhalation (4% induction, 2% maintenance) and ice block, respectively. The head of the anesthetized mouse was fixed in a stereotaxic frame using a nose cone and ear bars. Eyes were moistened with water-based lubricant, and the fur was cleared over the head with an electric shaver. A sterile cotton tip soaked in betadine was used to swab and clean the surface of the incision site. A midline incision was made on the scalp, and a small hole was drilled into the skull to expose the brain surface at the following stereotaxic coordinates selected to target the striatum, relative to bregma (anterior-posterior–1.00 mm, medial-lateral–1.80 mm). Cells were gently drawn up into a blunt-tipped 26G needle attached to a 10μl syringe (Hamilton Company, Reno, NV). The syringe was then placed into a microinjection pump (UltraMicroPump III, World Precision Instruments, Sarasota, FL) attached to the stereotaxic frame and lowered slowly into the injection site at a depth of 3 mm for adult mice and 1 mm for postnatal mice. The microinjection pump controlled the injection of cells at a flow rate of 0.5μl/min, after which the needle was left in place for 5 mins to ensure complete diffusion of cells and avoid backflow. Following the slow withdrawal of the needle, the hole was closed with dental cement, the incision was sutured, and the mice were observed until full recovery from anesthesia on a heated pad. Mice were monitored weekly. Animals assessed for survival were monitored until they lost more than 15% of their original body weight and/or had clinical symptoms of tumor development (hydrocephalus, lethargy, hunched posture). BRAF-2341 and BRAF-M34 displayed characteristics of HGG, such as hypercellular proliferation of atypical glial cells, focal necrosis, high microvascular proliferation, and occasional mitotic figures. RCAS-BRAF showed consistent and invasive growth into various brain regions, including the ipsilateral hippocampus, thalamus, white matter, lateral septal complex, somatomotor isocortex, and contralateral hemisphere **(Figure S2I)**. Histologically, RCAS-BRAF tumors exhibited hypercellular proliferation of malignant ovoid to spindled cells arranged in fascicles and sheets, with lower necrosis and microvascular proliferation compared to BRAF-2341 and BRAF-M34.

**Drug Preparation and Treatment**

Dabrafenib (cat#S2807; Selleckchem, Houston, TX) and Trametinib (cat#S2673; Selleckchem) were dissolved in DMSO at a concentration of 10 mg/ml and 2 mg/ml, respectively, and stored at -20°C. For *in vivo* experiments, a fresh solution of the drug was prepared on each treatment day, by thawing the stock solution in a warm water bath and diluting it in an aqueous mix of 0.5% hydroxypropyl methylcellulose (HPMC) and 0.2% tween solution to a final concentration of 30 mg/kg dabrafenib and 1 mg/kg trametinib. Drugs were delivered by oral gavage once daily. For survival analysis, dabrafenib was given at 30mg/kg daily by oral gavage. Trametinib was given at 1mg/kg daily by oral gavage. BioXCell anti-PD-L1 (clone 10F.9G2; ThermoFisher Scientific) was given at 20 mg/kg i.p. at day 1 and 8 mg/kg every 2 days thereafter, BioXCell anti-CTLA-4 (clone 9D9; ThermoFisher Scientific) was given at 8 mg/kg i.p. at day 1 and 2 mg/kg every 2 days thereafter.

For *in vitro* experiments, stock solutions of dabrafenib and trametinib were dissolved at 10 mM concentration in DMSO (3176-100 ml; ThermoFisher Scientific), aliquoted, and stored at -80 $^{o}$C. For measuring viability and constructing dose-response curves, cells were dissociated with Accutase (NC9389010; ThermoFisher Scientific) and $10^4$ cells were plated per well in a 96-well half-white plates (3688; ThermoFisher Scientific) using the corresponding vessel. After 24 hours, a serial dilution of 10μM dabrafenib or trametinib was added in triplicate to the seeded cells. After 72 hours, the cell viability was measured with CellTiter-Glo Luminescent Cell Viability Assay (G7570; Promega, Madison, WI). The percentage of surviving cells (normalized to DMSO control) was plotted as a function of log10 drug concentration. The sigmoidal curves were generated with GraphPad Prism software (GraphPad Prism version 10.0.0, GraphPad Software, Boston, Massachusetts USA, www.graphpad.com), whereby the IC50 concentrations were calculated.

**T cell depletion**

For survival study using T cell-depleted tumor-bearing mice orthotopically implanted with BRAF-M34 glioma cells, 200μg of anti-CD4 (clone GK1.5, Bio X Cell) and anti-CD8a (clone 53-6.7, Bio X Cell) were administered via intraperitoneal injections 3 days before tumor cell implantation and then every 4 days thereafter until the survival endpoint.

**Magnetic Resonance Imaging**

All MR studies were performed at 7T Bruker BioSpec system (Bruker, Billeria, MA) equipped with BGA-12S gradient insert and interfaced to ParaVision 360 V. 3.5. A 4-element receive-only mouse CryoProbe™ and 86mm volume coil were used for MR imaging. Gadobutrol solution (Gadavist; diluted 1:10 with saline; Bayer, Whippany, NJ) was administered intraperitoneally before MRI. Animals were anesthetized with 2.5% isoflurane mixed in air and $O_2$ and maintained with 1-1.5% isoflurane during the imaging procedure. Core body temperature was maintained at 37°C using a warm water circulator pad and a temperature controller. The animal's respiration was also monitored via a pneumatic pad placed under the animal (SA Instruments, Inc., Stony Brook, NY). T2w Turbo RARE sequence was used to obtain coronal (TR/TE = 2000/30 ms, FOV = 20 x 20 $mm^2$, matrix = 256 x 256, slice thickness = 0.5 mm, RARE Factor = 4, NEX = 2, scan time = 4 min 16 sec) and transverse (TR/TE = 2500/30 ms, FOV = 20 x 20 $mm^2$, matrix = 256 x 256, slice thickness = 0.5 mm, RARE Factor = 4, NEX = 2, scan time = 5 min 20 sec) structural MRI.

**Tissue Collection and Processing of Mouse Samples**

Mice were intraperitoneally injected with ketamine (100mg/kg) and xylazine (10mg/kg) to induce deep anesthesia, and subsequently transcardiac perfusion was performed first with phosphate-buffered saline followed by 4% paraformaldehyde (PFA)/PBS or 10% neutral buffered formalin. Brains perfused with 4% PFA were then resected, post-fixed in 4% PFA/PBS for 2 hours on ice, cryoprotected in 15% sucrose/PBS for 24 hours and then 30% sucrose/PBS for at least 48 hours, followed by embedding in Tissue-Tek O.C.T. Compound (Sakura Finetek USA, Inc., Torrance, CA). The embedded tissues were stored at -80 °C until sectioned. 10μm-thick coronal sections of the brain were cut on a cryostat (Leica, Wetzlar, Germany), collected onto Superfrost Plus slides (Menzel Glaser, VWR) and air dried for 1h before storing at -80°C until stained. Brains perfused with 10% formalin were post-fixed in 10% formalin for 24 hours at room temperature, then dehydrated in 70% ethanol. They were then embedded in paraffin and sectioned using Histo-Tech protocols (Histo-Tec Laboratory Inc., Hayward, CA; http://histoteclab.com/).

**Immunofluorescence Staining on Frozen Tissue Sections of Mouse and Human Samples**

Frozen 10μm sections on Fisherbrand™ Superfrost™ Plus Microscope Slides (ThermoFisher Scientific) were washed briefly in PBS and blocked with PBS containing 0.3% Triton X-100, 5~10% normal donkey serum, and/or normal goat serum for 1 hr at room temperature (RT). The sections were then incubated with primary antibodies at RT overnight in a humidified chamber, washed briefly in PBS three times next day, followed by 1h incubation at RT with secondary antibodies, the slides were then washed three times with PBS, and cover slipped with a DAPI-containing mounting solution (VectaShield H20002; Vector Laboratories, Newark, CA) and subjected to fluorescence and confocal microscopic analysis. For multiplex immunofluorescence, some primary antibodies were incubated simultaneously. The following primary antibodies were used: chicken anti-Nestin (ab134017; Abcam, Waltham, MA), chicken anti-GFP (600-901-215; Rockland Immunochemicals, Pottstown, PA), chicken anti-PLP1 (AB15454; Millipore Sigma, Burlington, MA), goat anti-PDGFRα (AF1062; R&D Systems, Minneapolis, MN), mouse anti-Cre recombinase (MAB3120; Millipore Sigma, Burlington, MA), mouse anti-BRAF V600E (ab228461; Abcam), mouse anti-NeuN (MAB377; Millipore Sigma, Burlington, MA), mouse anti-PD-L1 (329702; BioLegend, San Diego, CA), mouse anti-MHC I (17-5958-82; ThermoFisher Scientific), rabbit anti-CD4 (ab183685; Abcam), rabbit anti-CD68 (76437; Cell Signaling), rabbit anti-Iba1 (019-19741; Dako), rabbit anti-Olig2 (AB9610; Millipore Sigma, Burlington, MA), rabbit anti-Ki67 (ab15580; Abcam), rat anti-MHC II (NBP1-43312; Novus Biologicals), rat anti-CD133 (14-1331-82; ThermoFisher Scientific), rat anti-GFAP (13-0300; ThermoFisher Scientific), rat anti-MBP (ab7349; Abcam), rat anti-CD8a (50-112-9493; ThermoFisher Scientific), rat anti-PD-L1 (A14764; ThermoFisher Scientific), and rat anti-CD4 (100402; BioLegend). Secondary antibodies raised in donkey or goat and conjugated to Alexa Fluor 488, Alexa Fluor 555, Alexa Fluor 594, Alexa Fluor 647, or Alexa Fluor 790 were purchased from Jackson ImmunoResearch Labs (West Grove, PA) or Invitrogen (now ThermoFisher Scientific) and used at 1:200 dilution.

**Hematoxylin & Eosin Staining, Immunohistochemistry and Bulk RNA Sequencing**

Patient tissue was processed according to well-established histologic protocols, including fixation in 10% neutral buffered formalin, paraffin embedding, sectioning (5µm), and standard histochemical staining using hematoxylin and eosin (H&E). H&E-stained slides were scanned with a 20x objective on Aperio AT2 slide scanner (Leica Biosystems). Comprehensive diagnostic evaluation included a review by a board-certified pathologist specialized in anatomic pathology and neuropathology, as well as a senior fellow in neuropathology. Immunohistochemical staining of consecutive patient tumor sections was performed on a VENTANA Discovery ULTRA autostainer (Ventana). FFPE tissue sections were deparaffinized and rehydrated. Antigen retrieval was performed using CC1 buffer (05424569001; Roche) for 32 minutes at 91°C. Primary antibodies were applied at a dilution of 1:100 for 60 minutes: Anti-BRAF (mutated V600E) antibody clone VE1 (ab228461; Abcam), PD-L1 XP Rabbit mAb clone E1L3N (Cell Signaling, 13684). Detection was performed using the ChromoMap DAB Kit (05266645001; Roche). Slides were scanned with a 20x objective on the PhenoImager Fusion (Akoya Biosciences). Immunohistochemical staining of RCAS-BRAF mouse brain sections was performed by Histo-Tec. For bulk RNA sequencing, RNA extraction from paired FFPE-preserved specimens from patient 12, library preparation, and RNA sequencing was performed as previously described in Schreck et al., 2021.

**Differentiation Assays**

Dissociated cells were plated in Lab-Tek II CC$^2$ chamber slides (154941; ThermoFisher Scientific) coated with 10 ug/ml laminin or 0.01% poly-l-ornithine at 3 x $10^4$ cells per well. Two different growth conditions were used for this assay: medium without growth factors and medium supplemented with 60 nM triiodothyronine (T3; Millipore Sigma) and 10 ng/mL CNTF (Peprotech, Cranbury, NJ), termed as “differentiation” medium. Fresh medium and drugs (0.7μM dabrafenib and 0.3μM trametinib) was added every 3 days, to promote oligodendrocyte differentiation. The duration of the treatment was 13 days, and the experiment was performed in two technical replicates. Cells were fixed with 4 % PFA (EMD) for 10 min at RT and stained with primary antibodies overnight and secondary antibodies for 1 hour before being mounted in Vectashield mounting medium. Patient-derived glioma cell lines (STN49 and aGBM5) were seeded in MaTek chamber slides at high density (300,000 cells/well). After 24 hours of incubation, cells were treated either with 10 μg/ml bovine serum albumin (BSA, A841; Millipore Sigma) or 10μg/ml human recombinant galectin-3 (450-38; ThermoFisher Scientific) and incubated for 72 hours.

**nELISA Assay**

The high-plex secretome was profiled by NomicBio (https://nomic.bio/), which is offered as a service by the company. However, this platform does not detect mouse targets. For this study, secreted cytokines from the human cell lines (STN49 and aGBM5) were screened with the nELISA platform (https://doi.org/10.1101/2023.04.17.535914). Briefly, cells from STN49 and aGBM5 were dissociated using Accutase, and $10^4$ cells per well were seeded in 96-well plates (7200309; ThermoFisher Scientific). For this experiment, three different medium conditions were used (medium supplemented with/without growth factors and differentiation medium (see Differentiation Assays)). In addition, the medium was supplemented with dabrafenib and trametinib (0.7μM and 0.3μM, respectively) or DMSO (10μM) as a control. 50μl of medium from each culturing/drug combination was collected in triplicate on day 1, 3, 6, 9, and 13 post-therapy, and at the same time, fresh medium was added to the cells. Finally, the frozen medium containing cell-secreted cytokines and the control medium (freshly prepared without cell culturing) were submitted for nELISA measurements. Data was made available on the Nomic Data Portal (https://portal.nomic.bio/login), whereby the clustergram (Figure S9A) was downloaded from the custom data report. In this analysis (Figure S9A), TIMP1 protein was identified as a stably expressed marker, remaining unaffected by medium or drug conditions. As a result, TIMP1 levels were subsequently used for the normalization of other marker expressions.

Raw nELISA levels (pg/ml) of Galectin-3 were divided by the corresponding TIMP1 (pg/ml) levels. Raw nELISA signals of IL-23 and IL-32 were normalized by the corresponding TIMP1 expression signals. Time course response curves were created in GraphPad Prism software v.10 (GraphPad Prism version 10.0.0, GraphPad Software, Boston, Massachusetts USA, www.graphpad.com), and two-way ANOVA (without repeated measures) was used for statistical analysis. Data is represented as means ± SEM of three technical replicates, and significance is represented as *P<0.05, **P<0.01, ***P<0.001, and ****P<0.0001.

**Immunocytochemistry**

Prior to staining, cells were permeabilized with 0.25% Triton-X in PBS ($Ca^{2+}/Mg^{2+}$) for 5 minutes at room temperature, washed with PBS, and blocked with 10 % normal donkey serum (NC9624464; ThermoFisher Scientific) for 1 hour at room temperature on a gentle shaker. After this step, the blocking solution was aspirated and 150μl of primary antibody cocktail was added to each well: chicken anti-Nestin (ab134017; Abcam), goat anti-Olig2 (AF2418; R&D Systems), rat anti-mouse PD-L1 (A14764; ThermoFisher Scientific), mouse anti-human PD-L1 (329702; BioLegend), and/or rat anti-MBP (ab7349; Abcam). Cells in the Galectin-3 experiment were incubated with the following primary antibodies: mouse anti-human PD-L1, rat anti-MBP, and rabbit anti-OPALIN (ab121425; Abcam).

The primary antibodies were incubated overnight at 4°C on a gentle shaker. On the next day, the primary antibody solution was aspirated, and the wells were washed twice with PBS, which was followed by 150μl of secondary antibody staining solution: donkey anti-chicken-488, donkey anti-goat-647, goat anti-rat-488, goat anti-mouse-594, whereas donkey anti-mouse 488, donkey anti-rat 594, goat anti-rabbit 647 secondary cocktail was used for cells treated with Galectin-3/BSA . After 1 hour of incubation, cells were washed twice with PBS and mounted in Vectashield mounting medium containing DAPI.

**Confocal imaging and image processing**

Stained 10μm thick coronal sections and cultured cells in chamber slides were imaged by laser scanning confocal microscopy (Zeiss LSM980), which was used to detect up to five fluorophores by laser excitation at 405, 488, 561, 639, and 730 nm wavelengths. Confocal images were acquired using a 20X objective. Images were imported into FIJI/ImageJ image analysis software for quantification of cellular density and fluorescence intensity in the regions of interest. All analyses were performed in a blinded fashion.

### Flow Cytometry and MACS

Mouse brains with BRAF-2341 tumor cells were resected, and single-cell dissociates were obtained using 15 units/ml of Papain for 30 min at 37°C, and tissue was dissociated using a P1000 pipette, strained through a 70µm cell strainer over a 50 ml conical tube at room temperature. Cells were washed in 10 ml of ice-cold, sterile 1 x phosphate-buffered saline (PBS, Ca2+, Mg2+-free) and centrifuged at 1300 rpm for 10 min at 4°C. For flow cytometric analyses of CD133+ and Nestin+ cell frequency, cells were resuspended using PBS (Ca2+, Mg2+-free) + 0.2 mmol/L EDTA, resuspended in FACS buffer (PBS+EDTA+0.5% BSA) at 1E6 cells/100µl, and incubated with primary antibodies, which were chicken anti-Nestin (1:100; ab134017; Abcam ) + goat anti-chicken-488 (1:500; ab150169; Abcam) and Prominin-1 biotin (1:100; 130-101-996; Miltenyi Biotec, San Jose, CA) + streptavidin-APC (1:500; 130-090-856; Miltenyi Biotec). Isotype IgG and single-channel antibodies were used as a control. Primary antibodies were incubated for 30 min on ice in the dark. Cells were washed twice in 1 ml of ice-cold 2% fetal bovine serum + PBS, secondary antibodies were added for 15 min, washed, and cells were resuspended in a FACS buffer containing Hoechst and strained immediately prior to analyses. Cells were analyzed by flow cytometry using the BD FACSCalibur (BD Biosciences, Franklin Lakes, NJ). The gating strategy for analyzing CD133 and Nestin positive cells involved multiple steps, including initial quality control gating to remove debris and dead cells based on their size and granularity (FSC-A vs. SSC-A), doublet discrimination (FSC-A vs. FSC-H) to exclude cell doublets and clumps, live/dead discrimination (PI or DAPI), and specific gating for CD133 and Nestin positive cells based on unstained controls and isotype controls that can help set the appropriate threshold for positivity. Raw data was processed and analyzed with FlowJo software (10.7.1; BD Biosciences).

Magnetic sorting was performed according to manufacturer's instructions, with the modification that $10^7$ cells were resuspended in 60 ml of ice-cold MACS buffer solution, incubated in 20µl of FcR blocking reagent for 10 min at 2-8°C, 20µl of anti-A2B5 (130-123-953; Miltenyi Biotec) were added per $10^7$ cells, and incubated at 2-8°C for 15 min. Cells were washed twice and passed onto a LS column in an OctoMACS separator (Miltenyi Biotec). Columns were washed three times with 500 ml of MACS buffer and placed in a collection tube. A2B5+ cells were collected by firmly pushing the plunger into the column and immediately used for RNA extraction.

### Mass Cytometry and Analyses

FVB/N mice injected with BRAF-2341 glioma cells were treated as indicated for 2 weeks (Figure 6A). Tumor cells were isolated and subjected to mass cytometry (CyTOF) utilizing a 42-antibody panel (Weiss UCSF Mouse Neural Tumor CyTOF Antibody Panel) (Simonds, Erin F et al. "Deep immune profiling reveals targetable mechanisms of immune evasion in immune checkpoint inhibitor-refractory glioblastoma." *Journal for immunotherapy of cancer* vol. 9,6 (2021): e002181. doi:10.1136/jitc-2020-002181.) To isolate tumor cells, tumor-bearing mice were perfused with 5 ml Ca/Mg-free phosphate-buffered saline by transcardial perfusion. Brains were resected, and macroscopically visible tumor areas were cut from normal (non-infiltrated) brain tissue, and the tumor was further processed on ice using two # 11 scalpels in scissor-like motions into tumor pieces of 1 $mm^3$. Tumor fragments were dissociated into single cells with an enzymatic cocktail of collagenase IV (LS004209; Worthington), soybean trypsin inhibitor (LS003587; Worthington), and desoxyribonuclease I (LS002007, Worthington) at $37^0$C for up to 45 minutes. Single cells were subjected to red blood cell lysis using ACD lysis buffer (1049201; Gibco), following manufacturer's instructions. Viable cells were counted and stained with natural abundance cisplatin viability staining reagent (ALX-400-040; Enzo). Cells were frozen in 10% DMSO in 1e7 cells/ml aliquots in liquid nitrogen until ready for analyses.

1 ml of 1X Barcode Perm Buffer (Fluidigm) was added to isolated tumor cells, spun at 600x g for 5 minutes, vacuum aspirated to ~50µl and 750µl 1X Barcode Perm Buffer was added again to cells. 100µl of resuspended barcode stocks (two sets of 20-plex barcodes SCAC) were added to corresponding cell tubes and mixed by pipetting. Cells were incubated for 30 minutes at room temperature on RotoMix shaker at max speed, spun down, the excess buffer was aspirated, and cells were washed three times in cell staining media (CSM), and finally, 100µl CSM was added to each cell tube. For each barcode pool, barcoded cells were combined and strained into appropriate 5 ml FACS tubes and the volume of pooled tubes was brought up to 5 ml with CSM. Pooled barcoded cells were spun at 600x g, for 5 minutes, the supernatant was aspirated, and cells were vortexed to disrupt pellets and resuspended at 47µl per 1E6 cells. 2µl mouse Fc Block (clone 2.4G2; 0.5 mg/ml) per 1E6 cells were added to each tube, stained for 30 minutes at room temperature on RotoMix shaker at max speed. Cells were washed two times in 3 ml CSM, spun at 600x g for five minutes, excess CSM was aspirated, and the pellet was vortexed for 10 seconds to disrupt the pellet. Pellets were resuspended at 35µl per 1E6 cells. Surface antibody staining (Antibody Cocktail A) was performed first, with 15µl of Antibody Cocktail A per 1E6 cells added and mixed by pipet. Cells stained for 30 minutes at room temperature on RotoMix shaker at max speed and washed twice in 3 ml CSM, spun down at 600x g for 5 minutes, excess liquid aspirated, and pellet disrupted by 10-second vortex. Intracellular antibody staining (Antibody Cocktail B) was performed next by adding 3 ml Perm-S Buffer (Fluidigm) to the disrupted cell pellet and incubating

it on ice for 30 minutes. Barcoded cells were spun at 600x g for 5 minutes and supernatant vacuum aspirated. Cell pellets were resuspended with Perm-S Buffer (Fluidigm) at 35µl per 1E6 cells and mixed by pipet. 15µl of Antibody Cocktail B was added per 1E6 cells, mixed by pipet, and stained for 30 minutes at room temperature on RotoMix shaker at max speed. Cells were washed first with 3 ml Perm-S Buffer (Fluidigm), spun at 600x g for 5 minutes, supernatant aspirated, and pellet vortexed for 10 seconds and washed a second time with 3 ml CSM followed by spin, aspiration, and vortex. Cells were then resuspended in 500µl CSM and transferred to 15 ml polypropylene conical tubes and treated with 4.5 ml Ir-intercalator cocktail. Cells were stored at 4$^{o}$C overnight.

To prepare samples for CyTOF, 9 ml CSM was added per tube, cells were counted, then spun at 600x g for 5 minutes and vacuum aspiration left ~100 µl residual volume. To each tube, 800µl MaxPar PBS (Fluidigm) was added, mixed, and transferred to an Eppendorf Protein Lo-bind 1.5 ml tube, spun at 600x g for 5 minutes, then vacuum aspirated to ~100 µl residual volume. Cells were washed twice with 1.4 ml MilliQwater, spun at 600x g for 5 minutes and vacuum aspirated to ~100 µl residual volume. The barcoded cells were then resuspended in 1 ml 1X Four-Element EQ beads (Fluidigm), strained through blue-cap FACS tubes, and cells were recounted. CyTOF was performed on 7.87E6 cells in Pool 1 and 1.23E7 cells in Pool 2. FCS files were concatenated into a single FCS file utilizing the FCS concatenation tool from Cytobank.

Data was normalized using the Premessa normalizer tool in R (https://github.com/ParkerICI/premessa). CyTOF data was debarcoded and the panels were renamed using the same Premessa tool as above in R. The normalized, debarcoded, and renamed FCS files were uploaded to Cytobank for further downstream analyses of the data, including isolation and clustering of CD45+ cells with Phenograph. The fcs file containing the CD45+ pool with Phenograph coordinates was downloaded from Cytobank and loaded to the Cytofkit2 ShinyAPP via R (https://github.com/JinmiaoChenLab/cytofkit2), whereby the clusters were annotated and visualized on heatmaps and t-SNE plots.

**cDNA Synthesis and Quantitative PCR**

cDNA synthesis was performed using the iScript cDNA Synthesis Kit (Biorad). Quantitative RT-PCR was run using SYBR Green PCR Master Mix (ThermoFisher Scientific) using a cycling program of 10 min at 95$^{o}$C followed by 40 cycles of 95$^{o}$C for 15 sec and 95$^{o}$C for 60 sec. All primer sequences are available in Table S4.

**RNA Sequencing and Analysis**

RNA concentration was quantified by Nanodrop (Nanodrop Technologies, Wilmington, DE) and 0.4 µg of each sample was submitted to Novogene (Novogene Corporation Inc., Sacramento, CA) for RNA sample QC, library preparation and sequencing (Illumina NovaSeq6000, 20M paired reads). FASTQ files were uploaded to Galaxy (usegalaxy.org), an open-source, web-based platform for sequencing analysis where we performed quality control (FastQC), mapped the reads to hg19 or mm10 reference genome (STAR aligner), counted reads per annotated gene (FeatureCounts), and determined differentially expressed features (DESeq2). Pathway enrichment analysis was performed using gene set enrichment analysis (GSEA) combined with pathway definitions from MSigDB, KEGG, and Reactome databases (clusterProfiler). GSEA was run using pre-ranked mode for all genes based on log2 fold change derived from the differential expression analysis. Pathways with a normalized enrichment score >1 or <-1 at FDR <10% were considered significant. Heatmaps were generated using the ComplexHeatmap package in R.

**Single-cell Library Preparation and Sequencing**

C57BL/6 mice injected with RCAS-BRAF glioma cells were treated as indicated for 2 weeks. Brains collected after intracardiac perfusion with 20ml of PBS were dissociated into single-cell suspensions using the mouse tumor dissociation kit (130-096-730; Miltenyi Biotec) and the gentleMACS Octo Dissociator according to the manufacturer's protocol. Single-cell libraries of dissociated single cells were prepared using the BD Rhapsody™ platform (BD Biosciences). Single-cell suspensions obtained from the brains of individual tumor-bearing mice were stained with oligonucleotide-conjugated Sample Tags using the mouse Single-Cell Multiplexing Kit (cat# 633793) following the manufacturer's protocol (BD Biosciences). Sorted live cells from barcoded samples were counted and pooled at equal ratios and loaded onto a BD Rhapsody cartridge for capture. Cell capture and library preparation were completed using the BD Rhapsody Whole Transcriptome Analysis (WTA) Amplification Kit (cat# 633801). Briefly, cells were captured with beads in microwell plate, followed by cell lysis, bead retrieval, cDNA synthesis, template switching and Klenow extension, and library preparation following the BD Rhapsody protocol. The quality of the final libraries was assessed by using an Agilent's Bioanalyzer high sensitivity analysis and concentrations were measured by a Qubit method. The final libraries were diluted and normalized to 4nM and then pooled for paired-end sequencing on NovaSeq 6000 sequencer (Illuminia, San Diego, CA) in Novogene (Novogene Corporation Inc., Sacramento, CA).

**Analysis of Single-cell RNA Sequencing Data**

Rhapsody data were processed using the Seven Bridges Genomics cloud platform (San Francisco, CA) and BD Rhapsody Sequence Analysis Pipeline (v2.0) to quantify transcript levels with default parameters against the GRCm38 genome for BD Rhapsody scRNA-seq data. The plug-in Lex BDSMK was run to separate the Sample Tags. After analysis, output files were imported into the R package Seurat (version 4) for downstream clustering analysis and visualization. Briefly, cells were filtered, normalized, and the top 2000 highly variable features and the top 20 principal components were selected for further clustering analysis. Fourteen clusters were identified with the default resolution, including tumor and immune cell clusters based on the expression of their specific markers. Gene Ontology (GO) pathway enrichment analysis for differentially expressed genes between groups was performed using the clusterProfiler package. For sub-clustering analysis of tumor cells, CD4 and CD8 T cells, TAMs/microglia and dendritic cells, cells from the clusters of interest identified in primary clustering were extracted, reclustered and annotated based on subtype marker genes. Heatmaps for genes of interest were generated using the ComplexHeatmap package.

**Statistical Analysis**

All significance calculations were conducted using GraphPad Prism software. Statistical significance was determined using an unpaired, two-tailed Student's t-test or two-way ANOVA method when more than two groups were being compared. Survival was analyzed by the Kaplan-Meier method using log-rank (Mantel-Cox) test. Quantitative data are reported as mean ± SEM, and any values of $P<0.05$ were considered statistically significant. *$P<0.05$, **$P<0.01$, ***$P<0.001$ and ****$P<0.0001$.
